# Mutational resilience of antiviral restriction favors primate TRIM5α in host-virus evolutionary arms races

**DOI:** 10.1101/2020.06.12.149088

**Authors:** Jeannette L. Tenthorey, Candice Young, Afeez Sodeinde, Michael Emerman, Harmit S. Malik

## Abstract

Host antiviral proteins engage in evolutionary arms races with viruses, in which both sides rapidly evolve at interaction interfaces to gain or evade immune defense. For example, primate TRIM5α uses its rapidly evolving “v1” loop to bind retroviral capsids, and single mutations in this loop can dramatically improve retroviral restriction. However, it is unknown whether such gains of viral restriction are rare, or if they incur loss of pre-existing function against other viruses. Using deep mutational scanning, we comprehensively measured how single mutations in the TRIM5α v1 loop affect restriction of divergent retroviruses. Unexpectedly, we found that the majority of mutations increase antiviral function. Moreover, most random mutations do not disrupt potent viral restriction, even when it is newly acquired via single adaptive substitutions. Our results indicate that TRIM5α’s adaptive landscape is remarkably broad and mutationally resilient, maximizing its chances of success in evolutionary arms races with retroviruses.

## INTRODUCTION

Mammalian genomes combat the persistent threat of viruses by encoding a battery of cell-intrinsic antiviral proteins, termed restriction factors, that recognize and inhibit viral replication within host cells. The potency of restriction factors places selective pressure on viruses to evade recognition in order to complete replication (Duggal & Emerman, 2012). In turn, viral escape spurs adaptation of restriction factors, by selecting for variants that re-establish viral recognition and thereby restriction (McCarthy, Kirmaier, Autissier, & Johnson, 2015). Mutual antagonism between viruses and their hosts thus drives cycles of recurrent adaptation, in prey-predator-like genetic arms races (Van Valen, 1973). These arms races result in the rapid evolution of restriction factors, which accumulate amino acid mutations at their virus-binding interface at a higher than expected rate (M. D. Daugherty & Malik, 2012).

Numerous restriction factors, including TRIM5α (Sawyer, Wu, Emerman, & Malik, 2005), APOBEC3G (Sawyer, Emerman, & Malik, 2004), and MxA (Mitchell et al., 2012), evolve rapidly as a result of arms races with target viruses. The resulting divergence between restriction factor orthologs can result in cross-species barriers to viral infection (Compton & Emerman, 2013; Kirmaier et al., 2010). Such barriers led to the initial identification of TRIM5α, during a screen for proteins that prevented HIV-1 (human immunodeficiency virus) from efficiently replicating in rhesus macaque cells (Stremlau et al., 2004). Rhesus TRIM5α could potently restrict HIV-1, whereas the virus almost completely escapes TRIM5α-mediated inhibition in its human host. Subsequent studies revealed that restriction of SIVs (simian immunodeficiency viruses) also varies across TRIM5α orthologs and that SIVs likely drove the rapid evolution of TRIM5α in Old World monkeys (McCarthy et al., 2015; F. Wu et al., 2013).

TRIM5α disrupts retroviral replication early in infection by binding to the capsid core of retroviruses entering the cell (Y.-L. Li et al., 2016; Maillard, Reynard, Serhan, Turelli, & Trono, 2007; Owens, Yang, Gottlinger, & Sodroski, 2003). Its binding causes the premature uncoating of the viral core (Stremlau et al., 2006), preventing delivery of the viral genome to the nucleus for integration. TRIM5α binds to the capsid via the unstructured v1 loop within its B30.2 domain (Biris et al., 2012; Sebastian & Luban, 2005). Experiments swapping the v1 loop between TRIM5α orthologs indicate that it is critical for recognition of capsid from many retroviruses (Ohkura, Yap, Sheldon, & Stoye, 2006; Perron, Stremlau, & Sodroski, 2006; Sawyer et al., 2005). In Old World monkeys and hominoids, rapid evolution of TRIM5α is concentrated within this v1 loop (Sawyer et al., 2005) (Figure 1A). Single amino acid mutations at these rapidly evolving sites can cause dramatic gains of restriction against HIV-1 and other retroviruses (Y. Li, Li, Stremlau, Lee, & Sodroski, 2006; Maillard et al., 2007; Yap, Nisole, & Stoye, 2005). However, it remains unclear whether such adaptive mutations are rare among all single mutational steps that might be randomly sampled during TRIM5α’s natural evolution.

**Figure 1.**
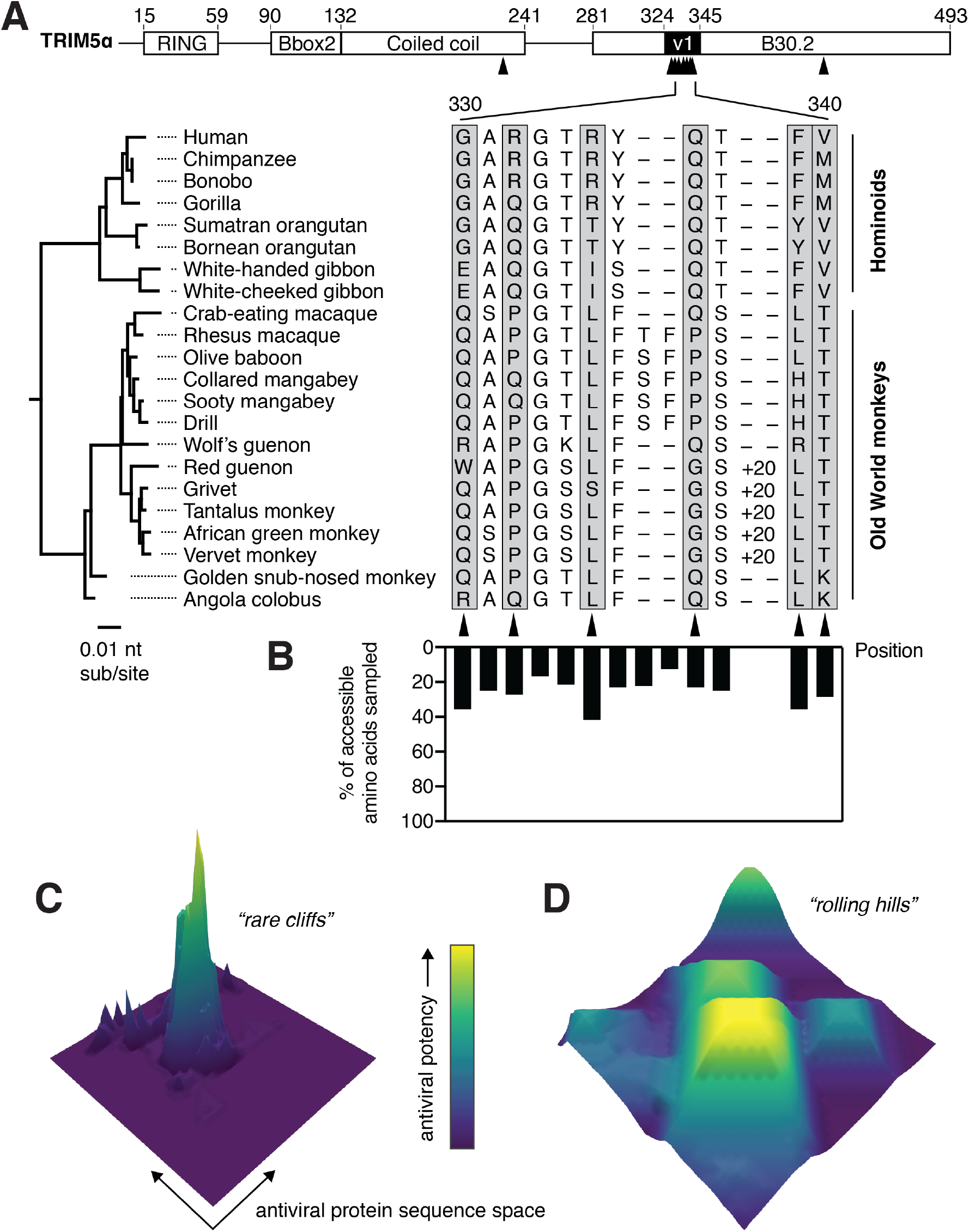
TRIM5α has sampled limited amino acid diversity, even at rapidly evolving positions. (**A**) Alignment of TRIM5α from simian primates. A 20-amino acid duplication in the v1 loop of the African green monkey clade is abbreviated as “+20”. Amino acid numbering follows human TRIM5α. Rapidly evolving residues are indicated with black arrows and gray boxes. (**B**) Evolutionarily accessible amino acids were defined as within 1 nucleotide of any codon in this alignment, and the fraction of accessible variants sampled among aligned sequences was determined for each position. (**C-D**) Theoretical possibilities for antiviral protein evolutionary landscapes, with antiviral potency represented in z and color axes as it varies with single point mutations. Fitness landscapes might be highly constrained (**C**) or permissive (**D**).

The functional consequence of all single mutations from a given protein sequence can be visualized as an evolutionary landscape (Smith, 1970), in which mutations are either beneficial (fitness peak), detrimental (fitness valley), or neutral. The topology of this evolutionary landscape, in terms of numbers of peaks and valleys and their connections, represents the adaptive potential of restriction factors in their evolutionary arms race with viruses. Previous studies that have empirically mapped evolutionary landscapes of conserved enzymes and transcription factors revealed that ligand-binding residues are highly intolerant to substitutions (Fowler et al., 2010; Guo, Choe, & Loeb, 2004; McLaughlin, Poelwijk, Raman, Gosal, & Ranganathan, 2012; Suckow et al., 1996). Moreover, mutations that allowed proteins to gain novel ligand specificity, even for closely related ligands, were rare among all possible substitutions (McLaughlin et al., 2012; Starr, Picton, & Thornton, 2017; Stiffler, Hekstra, & Ranganathan, 2015). In contrast, TRIM5α and other restriction factors can dramatically change antiviral potency via single mutations at viral interaction interfaces (M. D. Daugherty & Malik, 2012; Mitchell et al., 2012). However, since the frequency of such gain-of-function mutations is unknown, it is unclear whether virus-binding surfaces in rapidly evolving antiviral factors are subject to the same evolutionary constraints as previously mapped for other proteins.

Here, we investigated the adaptive landscape of antiviral specificity conferred by the rapidly evolving, capsid-binding v1 loop of TRIM5α. To our surprise, we found that, rather than the evolutionary landscape of TRIM5α being narrowly constrained among all possible amino acid substitutions, the majority of random mutations in the v1 loop resulted in gains of antiviral restriction. We found that the primary v1 loop determinant for TRIM5α’s restriction of HIV-1 and other lentiviruses is its net electrostatic charge. Furthermore, both rhesus and human TRIM5α proteins are highly resilient to mutation, in that they withstand more than half of all possible single amino acid mutations in the v1 loop without compromising their antiviral restriction abilities. This unexpectedly permissive landscape allows TRIM5α to sample a wide variety of mutations to maximize its chances of success in arms races with retroviruses.

## RESULTS

### A deep mutational scan of the TRIM5α v1 loop

Despite their rapid evolution, primate TRIM5α orthologs have sampled relatively limited amino acid diversity at rapidly evolving positions within the capsid-binding v1 loop (Figure 1A). For example, although single amino acid changes at residue 332 are responsible for dramatic differences in antiviral restriction (Y. Li et al., 2006), this residue repeatedly toggles between just three amino acids. The limited diversity is not due to evolutionary inaccessibility, since most amino acids that can be sampled with single nucleotide changes are not observed among primate TRIM5α orthologs (Figure 1B). There are two alternative explanations for this restricted diversity. First, it might suggest that adaptive gain-of-function mutations in TRIM5α are rare, with TRIM5α’s evolutionary landscape mainly consisting of fitness valleys with only a few mutational avenues to reach fitness peaks (Figure 1C). Conversely, the limited diversity might be a consequence of epistatic interactions with other sites that constrain amino acid sampling, or even simple chance. Under this scenario, TRIM5α’s evolutionary landscape may consist of numerous, wide peaks that tolerate substantial mutational variation (Figure 1D). We sought to differentiate between these possibilities by experimentally defining the evolutionary landscape of antiviral restriction, by both human and rhesus TRIM5α, over all possible single mutational steps in the v1 loop.

We took a deep mutational scanning (DMS) approach (Fowler et al., 2010) to measure the effect on antiviral restriction of all v1 loop single mutations in a pooled assay. We first generated a library of all single amino acid variants (including stop codons) within the rapidly evolving portion of the v1 loop (amino acids 330 to 340, Figure 1A), with a library diversity of 231 amino acid (352 nucleotide) variants (Figure 2A). The resulting TRIM5α variants were stably expressed via transduction into CRFK (cat renal fibroblast) cells, which naturally lack TRIM5α (McEwan et al., 2009). We transduced CRFK cells at a low dose to limit the integration of multiple variants into individual cells, thus generating a pool of cells each expressing a single TRIM5α point mutant. Libraries were represented with at least 500-fold coverage through all experimental steps to avoid bottlenecking library diversity.

**Figure 2.**
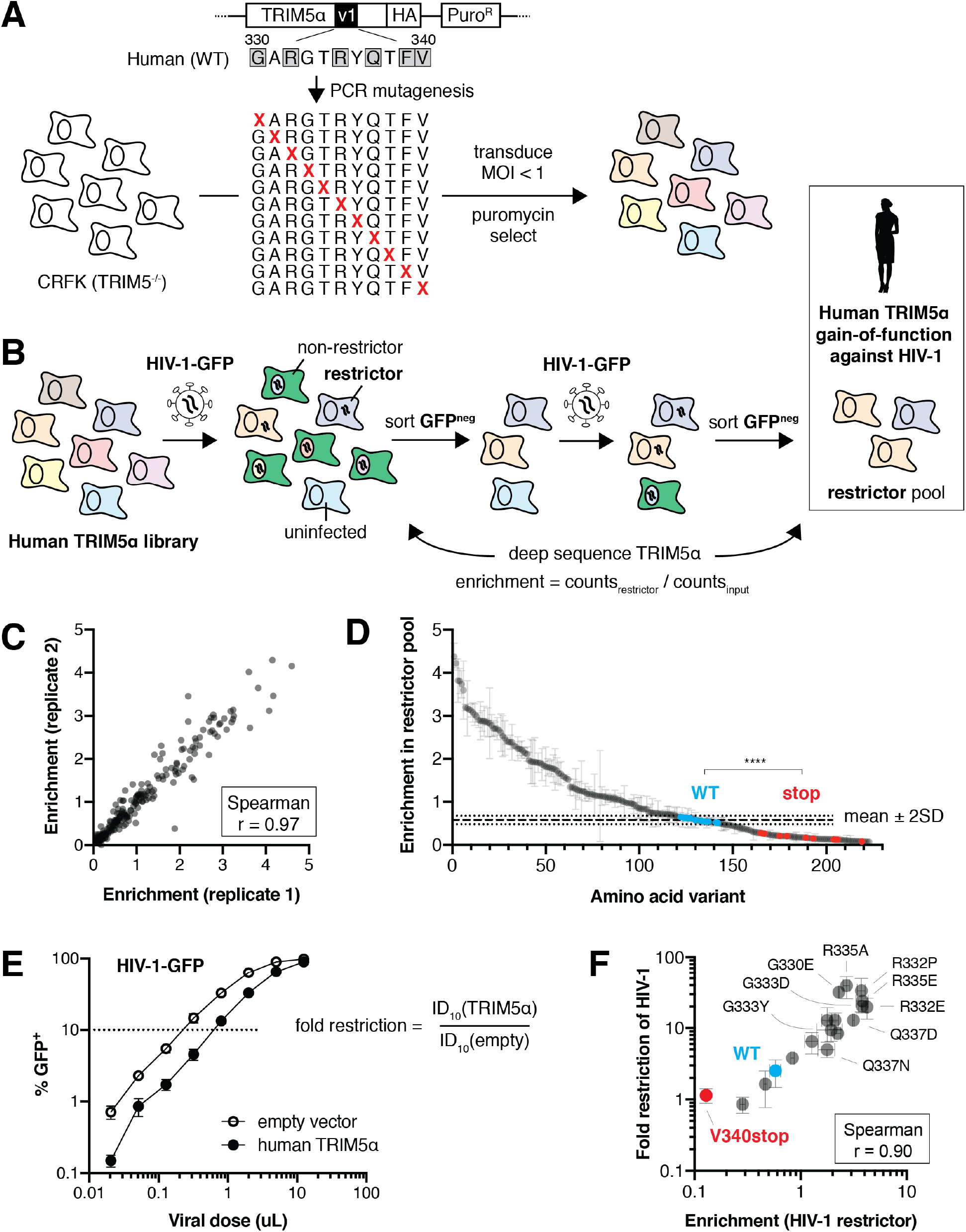
Selection scheme to identify human TRIM5α variants that gain HIV-1 restriction. (**A**) A DMS library, encoding all single amino acid variants within the v1 loop (rapidly evolving sites are boxed), was generated by PCR with degenerate NNS codons. The library was transduced into naturally TRIM5α-deficient CRFK cells at low MOI and selected using puromycin. Colors represent different TRIM5α variants. (**B**) Pooled TRIM5α-expressing cells were infected with HIV-1-GFP virus-like particles (VLPs) at a high dose. GFP-negative cells were FACS sorted, re-infected, and re-sorted. Restrictive TRIM5α variants were then sequenced, and variant frequencies were normalized to input representation. (**C**) Amino acid enrichment scores are highly correlated across 2 biological replicates. Each dot represents a unique amino acid sequence, averaged across synonymous codons. (**D**) Nonsense variants (red) are depleted relative to WT (blue) and most missense (gray) variants; ****p < 0.0001, student’s unpaired t-test with Welch’s correction. Enrichment is averaged across synonymous codons and replicates, except for WT variants, which are averaged only across replicates to better visualize variance (WT mean ± 2 standard deviations is indicated). (**E**) HIV-1 fold-restriction by TRIM5α was measured by the increase in ID_10_ (viral dose at which 10% of cells are infected) relative to an empty vector control. (**F**) Enrichment scores are highly correlated with HIV-1 restriction for re-tested variants. (**D-F**) Error bars, SD.

Human TRIM5α only weakly restricts HIV-1 (Jimenez-Guardeño, Apolonia, Betancor, & Malim, 2019; OhAinle et al., 2018). However, single amino acid mutations in the v1 loop can substantially increase restriction activity (Y. Li et al., 2006; Pham, Bouchard, Grütter, & Berthoux, 2010; Pham et al., 2013). To comprehensively assess how many single mutation variants of human TRIM5α had increased activity against HIV-1, we first performed a gain-of-function screen. We challenged the library of human TRIM5α variant-expressing cells with HIV-1 bearing a GFP reporter, at a dose infecting 98% of cells (Figure 2B). Because GFP expression becomes detectable only after integration of the HIV-1 proviral genome, cells expressing TRIM5α variants that restrict HIV-1 infection remain GFP-negative. However, ~2% of cells that were uninfected by chance would also be GFP-negative. Therefore, to enrich for cells expressing *bona fide* restrictive TRIM5α variants, we sorted the GFP-negative cells from the first round of infection and subjected them to a second round of HIV-1-GFP infection and sorting. Following this second round of selection, we deep sequenced the TRIM5α variants in the GFP-negative cell population. We normalized the count of each variant to its representation in the pre-selection cell population to determine its enrichment score, which should reflect the relative antiviral function of each TRIM5α variant.

Enrichment scores were highly correlated between two independent biological replicates (Figure 2C, Spearman r = 0.97). Furthermore, mutants bearing premature stop codons, which should be non-functional and depleted from the restrictor pool, were all among the most depleted variants (Figure 2D, red). Despite the weak (~2-fold) restriction of HIV-1 by wildtype (WT) human TRIM5α, variants containing synonymous nucleotide changes (no amino acid changes compared to WT, blue) had significantly higher enrichment scores than those containing stop codons (p < 0.0001, student’s unpaired t-test with Welch’s correction), confirming that the assay worked as expected.

To investigate whether enrichment scores were truly representative of increased antiviral function, and to validate some of the novel amino acid changes that appeared to result in increased restriction, we made 16 targeted missense mutants from across the enrichment spectrum and challenged them individually with HIV-1-GFP. We determined their fold-restriction by determining the relative viral dose required to infect 10% of cells (ID_10_) expressing a TRIM5α variant compared to an empty vector control; a larger viral dose is required to overcome TRIM5α-mediated restriction (Figure 2E). We confirmed that several previously described gain-of-function variants (Y. Li et al., 2006; Pham, Bouchard, Grütter, & Berthoux, 2010; Pham et al., 2013) had increased antiviral activity and were highly enriched (Figure 2F: G330E, R332P, R332E, R335E). Moreover, we identified novel amino acid mutations that significantly increased antiviral activity, such as R335A and G333D, whereas moderately enriched variants (e.g., G333Y, Q337N) had correspondingly modest gains in HIV-1 restriction. Indeed, enrichment scores and fold-restriction were highly correlated across all mutants tested (Spearman r = 0.90). Thus, enrichment scores accurately reflect antiviral activity, validating our approach to simultaneously identify all single mutants with increased HIV-1 restriction. Therefore, in subsequent analyses, we use enrichment scores as a proxy for the antiviral restriction activity of TRIM5α mutants.

### Most single mutations in the v1 loop improve human TRIM5α restriction of HIV-1

Based on the limited amino acid diversity among primate TRIM5α v1 loops (Figure 1A-B), we expected that our DMS assay would reveal only a few beneficial mutations that improve human TRIM5α restriction of HIV-1. Contrary to this expectation, we found that more than half of all missense variants (115, 57%) had enrichment scores that fell more than two standard deviations above WT TRIM5α (Figure 2D). Even if we limited our analysis to amino acid variants that are evolutionarily accessible via single nucleotide changes from the WT *TRIM5α* sequence, this ratio did not change substantially (32, 54%). These enrichment scores represent dramatic gains in HIV-1 restriction, with the most potent variants (R332P and R335A) improving HIV-1 restriction ~15-fold relative to WT (33- and 39-fold restriction, respectively). Our findings indicate that the fitness landscape of the v1 loop is not narrowly constrained, but rather is remarkably permissive (Figure 1D), in that most single amino acid changes not seen in natural sequences enhance the ability of human TRIM5α to restrict HIV-1. Thus, TRIM5α has the capacity to readily evolve antiviral potency against HIV-1 via single mutations.

We analyzed whether a common biochemical mechanism could explain the unexpectedly high fraction of restrictive TRIM5α variants. We found that increased expression levels could explain some of the improvement in HIV-1 restriction, although several mutations (G333D, G333Y) improved restriction without increasing expression (Figure 3—figure supplement 1). In contrast, most gains in HIV-1 restriction could be completely accounted for by a reduction in the electrostatic charge of the v1 loop (Figure 3A), regardless of expression level. For example, mutation of the positively-charged residues 332 or 335 from the WT arginine (R) to any amino acid except lysine (K) significantly improved HIV-1 restriction (Figure 3A-B), consistent with previous reports on R332 variants (Y. Li et al., 2006). Mutation of uncharged sites to K or R decreased TRIM5α restriction of HIV-1, whereas introducing a negatively-charged aspartic acid (D) or glutamic acid (E) significantly increased HIV-1 restriction (Figure 3C). Since our DMS assay tests only one mutation at a time, all mutations to D or E occur in the context of at least one proximal positively-charged residue. Therefore, we infer that the position-independent benefit of introducing D or E derives from offsetting pre-existing positive charge in the v1 loop that is detrimental to HIV-1 restriction. Indeed, reducing the net charge of the v1 loop explains all of the highest enrichment scores (Figure 3D). Thus, we conclude that positive charge in the v1 loop is the dominant impediment to HIV-1 restriction by human TRIM5α.

**Figure 3.**
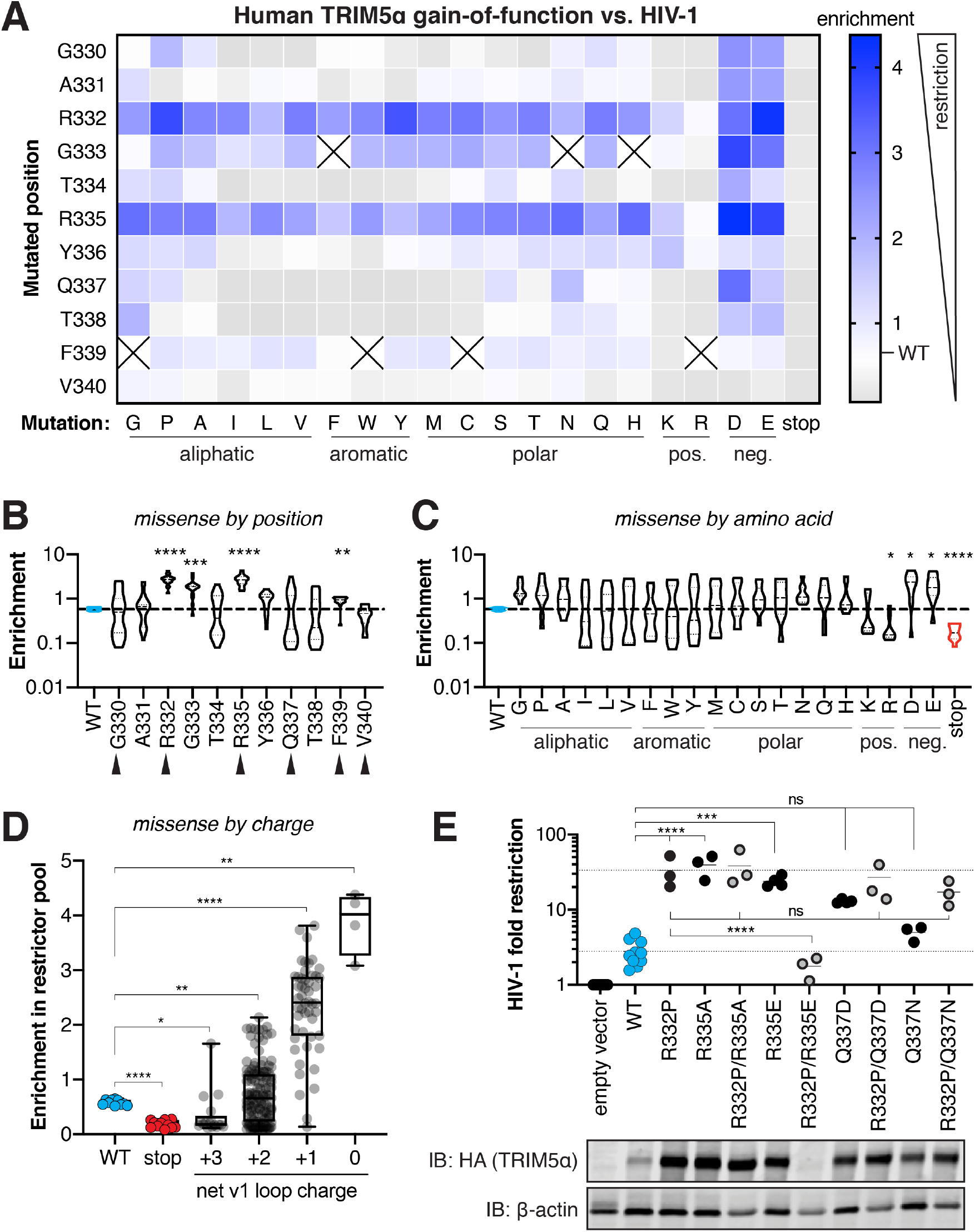
Many single mutations improve human TRIM5α restriction of HIV-1, primarily by removal of positive charge. (**A**) Enrichment in the HIV-1 restrictor pool relative to WT (white) for each TRIM5α variant, arrayed by position mutated and amino acid mutation, is indicated by color intensity. Variants marked with X were excluded due to low input representation. (**B-C**) Enrichment scores for each position across all amino acid variants (**B**) or each amino acid across all positions (**C**); statistics reported in comparison to WT. Rapidly evolving sites are indicated by black arrows. (**D**) Box plot of missense mutations grouped by their effect on the net v1 loop charge; WT has a net v1 charge of +2. (**E**) Gain-of-function mutations were tested against HIV-1 individually or in combination with R332P, and fold-restriction was determined as in Figure 2E. TRIM5α expression levels in CRFK cells was analyzed by immunoblot (IB) against the C-terminal HA tag. (**B-E**) *p < 0.05, **p < 0.01, ***p < 0.001, ****p < 0.0001; one-way ANOVA with Holm-Sidak’s correction for multiple comparisons and (**B-D**) correction for unequal variances.

**Figure 3–figure supplement 1.**
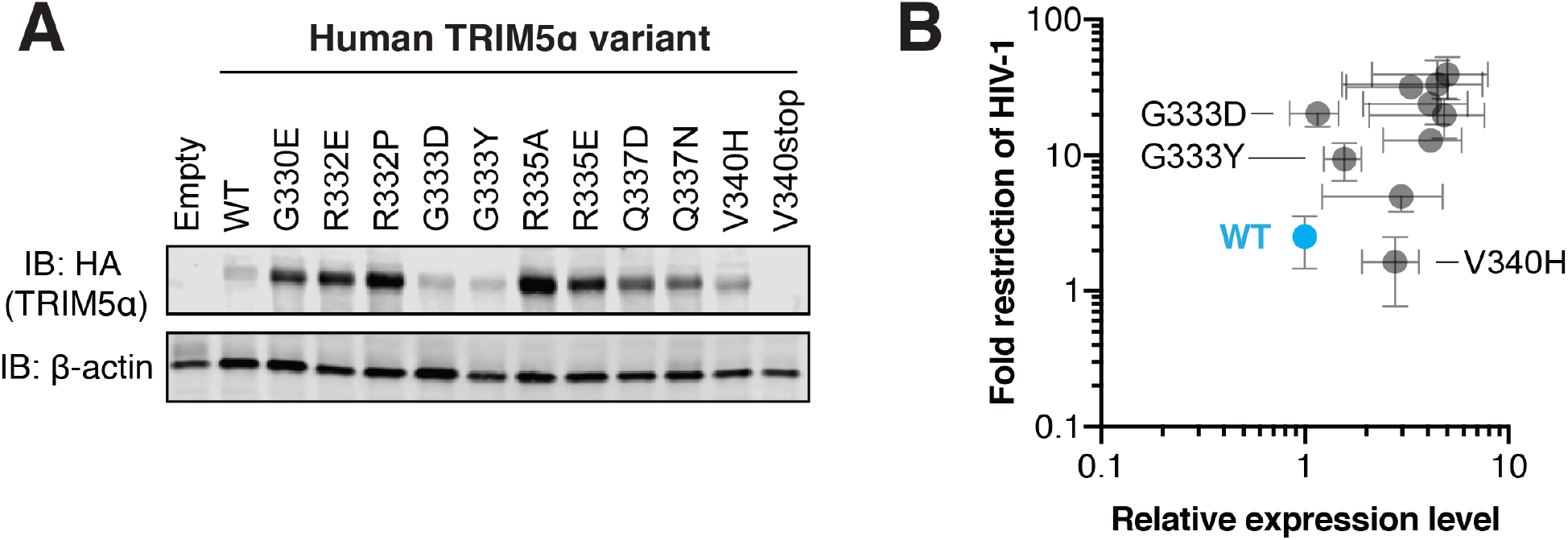
Some, but not all, human TRIM5α gain-of-function mutations against HIV-1 increase TRIM5α expression level. (**A**) Representative immunoblot (IB) for TRIM5α expression in CRFK cells. (**B**) TRIM5α-HA band intensity was normalized to β-actin, and then further normalized to WT TRIM5α to determine relative expression. Results from 3 independent experiments. HIV-1 restriction was calculated in Figure 2F. Error bars, SD.

Removal of positive charge, however, could not explain all of the improved HIV-1 restriction we observed. For example, despite its strict conservation in primate TRIM5α (Figure 1A), a glycine (G) at residue 333 compromises HIV-1 restriction. Mutation of G333 to most other amino acids significantly improves TRIM5α activity (Figure 3B). We confirmed this finding for several individual variants (Figure 2F: G333Y, G333D). We found a similar pattern for residue F339, which is disfavored for HIV-1 restriction, albeit not to the same extent as G333. Contrary to our initial expectations, there is only a weak association between rapidly evolving residues and residues whose mutation can significantly improve HIV-1 restriction: missense mutations in 3 of 6 rapidly evolving sites, versus 1 of 5 conserved sites, significantly improve HIV-1 restriction (Figure 3B). This result suggests that conserved positions in the vicinity of rapidly evolving sites possess unexpected potential to improve antiviral potency.

We also tested whether beneficial mutations might have additive effects on HIV-1 restriction by human TRIM5α. We combined several gain-of-function variants with the R332P mutation, previously described to potently restrict HIV-1 (Yap et al., 2005). However, we found that no double mutants tested increased HIV-1 restriction beyond that of R332P alone (Figure 3E). Instead, combination of one gain-of-function variant (R335E) with R332P resulted in loss of protein expression and HIV-1 restriction. Previous reports also found beneficial mutations to be either non-additive or interfering (Y. Li et al., 2006; Pham et al., 2010; 2013). Collectively, these results suggest that single mutations can confer most or all of the increased HIV-1 restriction potential onto human TRIM5α. Thus, remarkably, human TRIM5α appears to be located only one mutational step away from the numerous fitness peaks in its evolutionary landscape of adaptation against HIV-1.

Finally, we investigated whether gain-of-function mutations for HIV-1 restriction also conferred protection against other lentiviruses. We focused on lentiviruses that are v1 loop-dependent: either the entire v1 loop (Figure 4—figure supplement 1) or the R332P mutation from rhesus TRIM5α (Stremlau, Perron, Welikala, & Sodroski, 2005) could confer human TRIM5α with substantial antiviral function. In each case, WT human TRIM5α only weakly restricts these lentiviruses (Figure 4A). However, the charge-altering mutations R332P and R335A increased restriction of all lentiviruses we tested, including HIV-2, SIVcpz (SIV infecting chimpanzees), and SIVmac (SIV infecting rhesus macaques). Introduction of negative charge (R332E, R335E, G330E, G333D, and G337D) also selectively improved restriction of HIV-1, SIVcpz, and HIV-2 but not SIVmac. Thus, positive charge at positions 332 and 335 appears to be generally detrimental for lentiviral restriction. Furthermore, although TRIM5α fitness landscapes are lentivirus-specific, many of the mutations we tested increased restriction against other lentiviruses. These data suggest that the evolutionary landscape for lentiviral restriction by TRIM5α is likely to be generally permissive, as it is for HIV-1.

**Figure 4.**
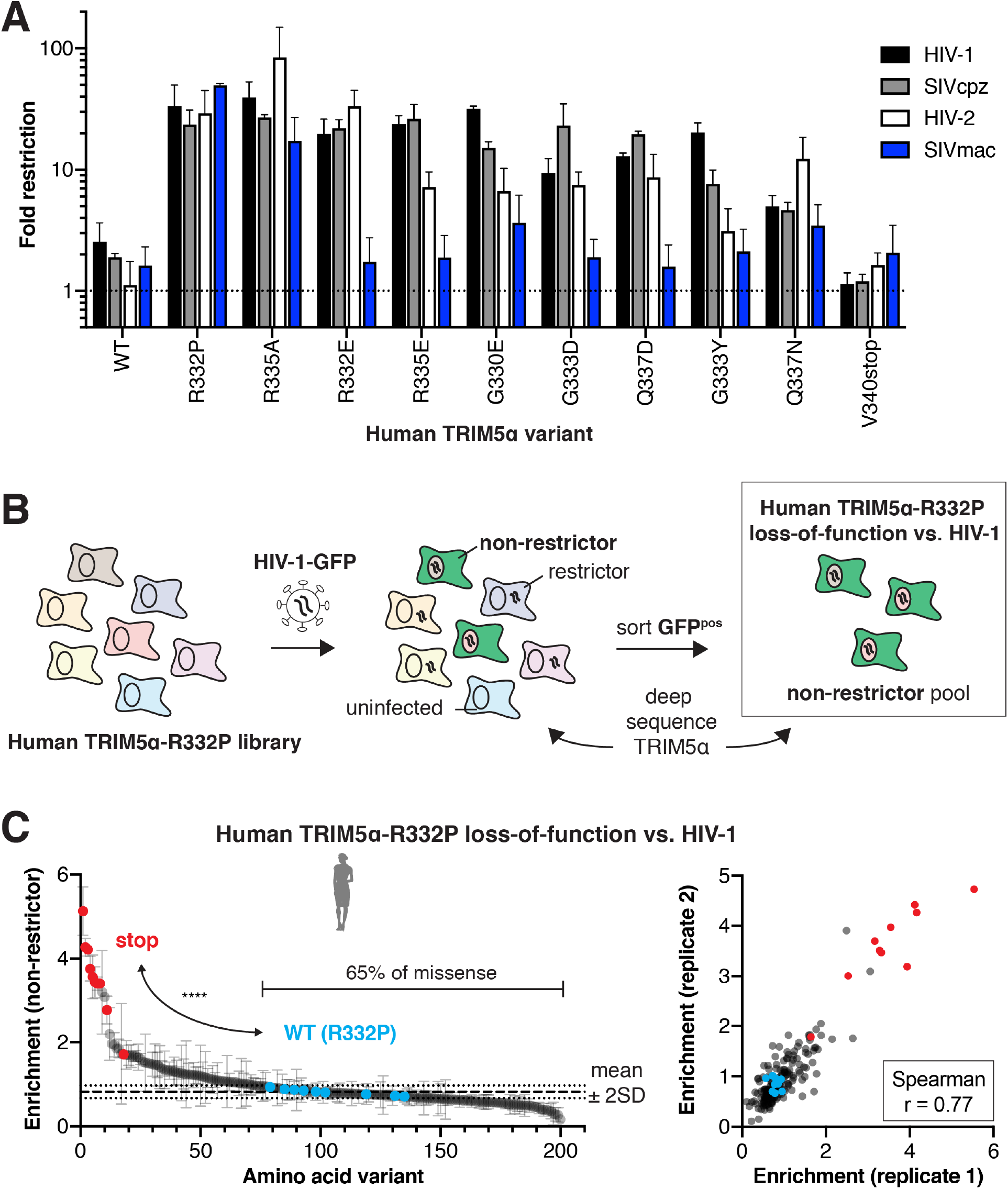
Evolutionary landscapes are generally permissive for evolving novel lentiviral restriction, which is resilient to most mutations once achieved. (**A**) CRFK cells expressing the indicated human TRIM5α variant were challenged with GFP-marked lentiviral VLPs to determine fold-restriction as in Figure 2E. (**B**) To determine whether newly acquired viral restriction tolerates mutations, a second human TRIM5α v1 DMS library was generated with R332P fixed in all variants. This library of cells was infected with HIV-1-GFP VLPs, and GFP-positive (non-restrictor) cells were sorted and sequenced. (**C**) Stop codon variants are highly enriched in the non-restrictor pool compared to WT (R332P, blue) variants (****p < 0.0001, student’s unpaired t-test with Welch’s correction), while 65% of all missense variants fall less than 2 SD above WT (R332P) mean. Enrichment scores between two biological replicates are well correlated. (**A, C**) Error bars, SD.

**Figure 4—figure supplement 1.**
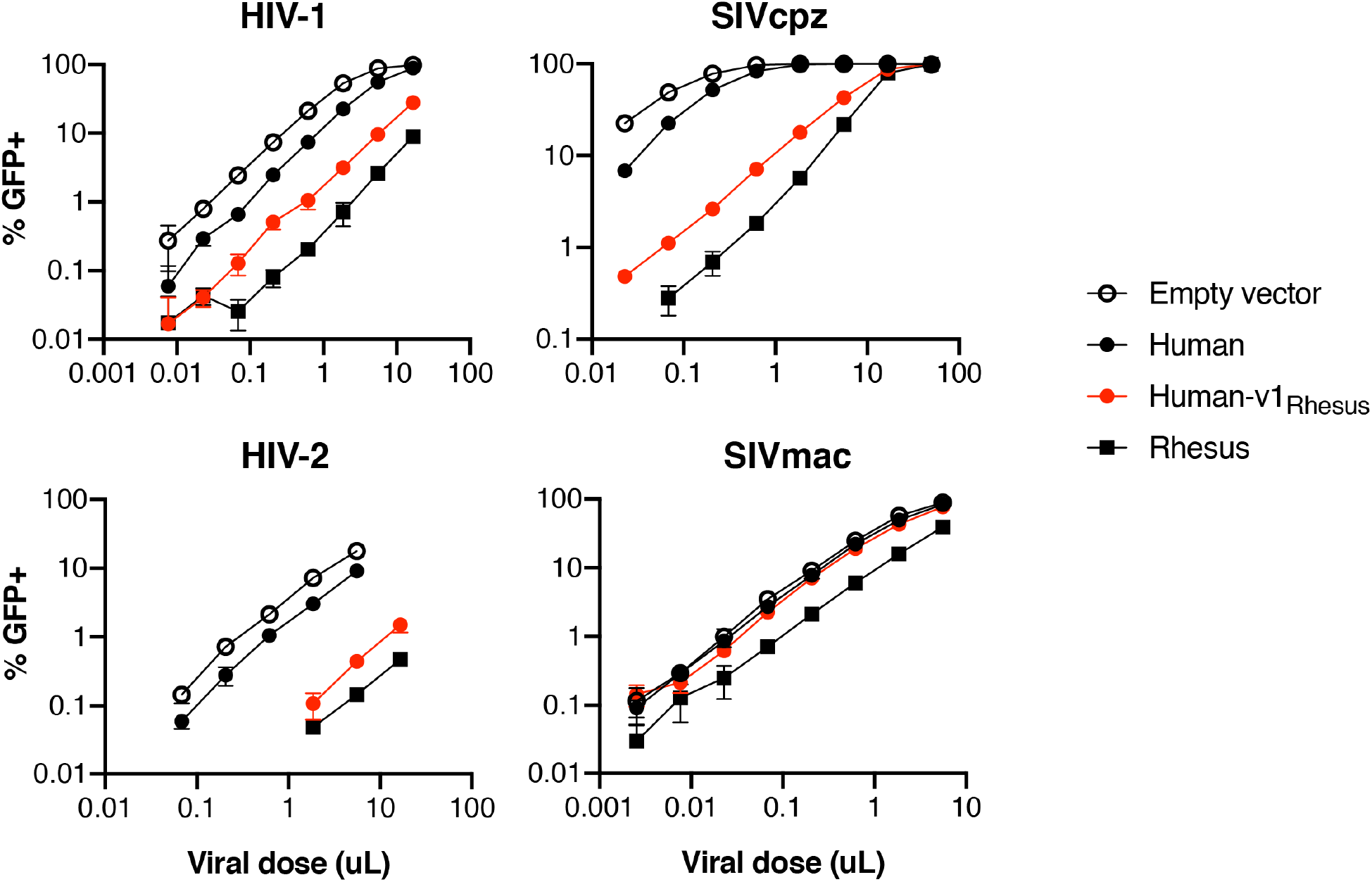
Lentiviral restriction by TRIM5α is v1-dependent. CRFK cells expressing human, rhesus, or human TRIM5α with the v1 loop exchanged for that of rhesus were challenged with GFP-marked lentiviral VLPs. Results representative of at least 3 independent experiments.

### TRIM5α restriction of HIV-1 is resilient to single mutations

Our data show that novel antiviral potency is readily attainable by single amino acid changes in human TRIM5α (Figures 2D, 4A). However, these gains might be just as easily lost through further mutation, since rapidly evolving antiviral proteins like TRIM5α continually adapt in their arms race with viruses. Therefore, in order to test whether newly acquired antiviral potency is fragile or resistant to mutation, we investigated the mutational resilience of the R332P variant of human TRIM5α, which inhibits HIV-1 ~15-fold more than WT (Figure 2F). To do so, we generated a v1 DMS library of human TRIM5α with R332P fixed in all variants. We challenged this pooled cell library with HIV-1-GFP, at a viral titer which human TRIM5α-R332P restricts to ~1% infection. In this case, we sorted and deep sequenced GFP-positive cells, so that enrichment (relative to initial representation) now reflects the degree to which each TRIM5α-R332P variant lost its antiviral function against HIV-1 (Figure 4B). As expected, we observed strong enrichment of stop codons in the non-restrictor pool and good correlation between biological replicates (Figure 4C).

As with WT human TRIM5α, addition of positive charge by mutations to K or R at most positions in the v1 loop reduced HIV-1 restriction (Figure 4—figure supplement 2). However, we found that the majority of missense variants (65%) did not, in fact, weaken HIV-1 restriction by the R332P variant of human TRIM5α. Thus, WT human TRIM5α is only one mutational step away from a fitness peak (Figure 3) that, once achieved, also exhibits a surprising degree of resilience to mutation. This implies that gains of restriction by TRIM5α are not likely to be compromised by its continued adaptation.

**Figure 4—figure supplement 2.**
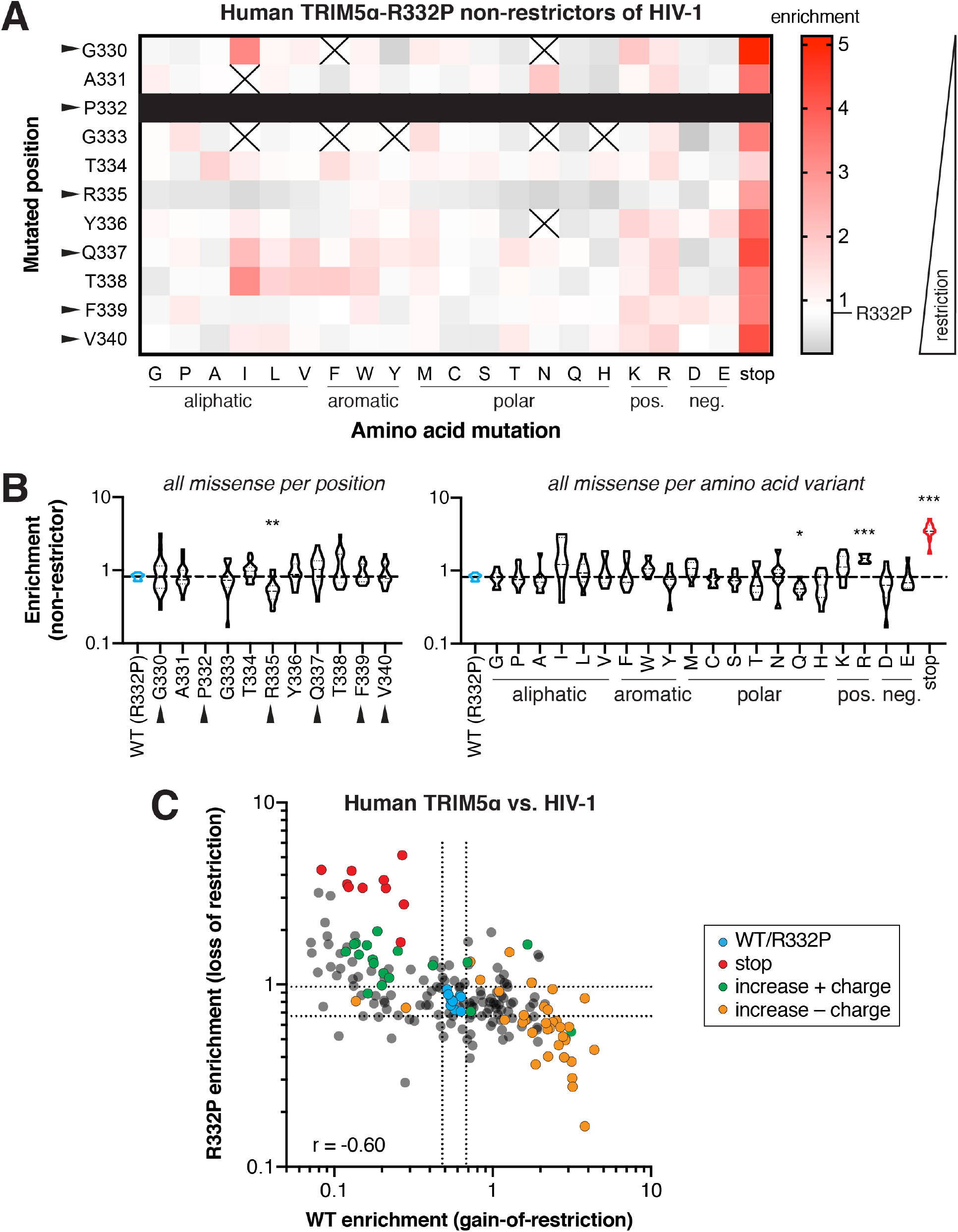
Biochemical preferences for human TRIM5α-R332P restriction of HIV-1. (**A**) Enrichment in the HIV-1 non-restrictor pool relative to R332P (white) for each human TRIM5α-R332P variant, arrayed by position and amino acid variant, is indicated by color intensity. Variants marked with X were excluded due to low input representation; no variation was present at position 332 (black box). Arrows indicate rapidly evolving sites. (**B**) Missense variants at each position (across all variants) or each amino acid (across all positions); one-way ANOVA compared to R332P with corrections for multiple comparisons (Holm-Sidak) and unequal variances (Geisser-Greenhouse); *p < 0.05, **p < 0.01, ***p < 0.001, ****p < 0.0001. (**C**) Positive charge is detrimental to human TRIM5α restriction of HIV-1, both for WT and R332P variants (Spearman r = −0.60). Missense mutations that increase positive charge (green) weaken R332P restriction, while those that negate positive charge (orange) improve WT restriction of HIV-1. Dashed lines indicate 2 SD above or below the mean enrichment for WT variants (blue) in each screen.

To determine if HIV-1 restriction is also resilient to mutation in a naturally-occurring TRIM5α variant, we next assessed the likelihood that random mutations disrupt viral restriction by WT rhesus macaque TRIM5α, which strongly restricts HIV-1 in a manner that strictly requires the v1 loop (Sawyer et al., 2005). Like with human TRIM5α, we constructed a library of cells each expressing a single rhesus TRIM5α variant, with each variant containing a single mutation in the v1 loop (note that the v1 loop is slightly longer in macaques compared to humans, Figure 1A). We then challenged this pool of cells with HIV-1-GFP and sorted GFP-positive cells for subsequent deep sequencing. Rhesus TRIM5α variants containing premature stop codons were strongly enriched in the non-restrictor pool, whereas WT variants were significantly depleted (Figure 5A). In contrast, half of all missense mutations (125, 51%) fell within two standard deviations of WT. Even missense mutations accessible by single nucleotide changes reflected this pattern (40, 55%).

**Figure 5.**
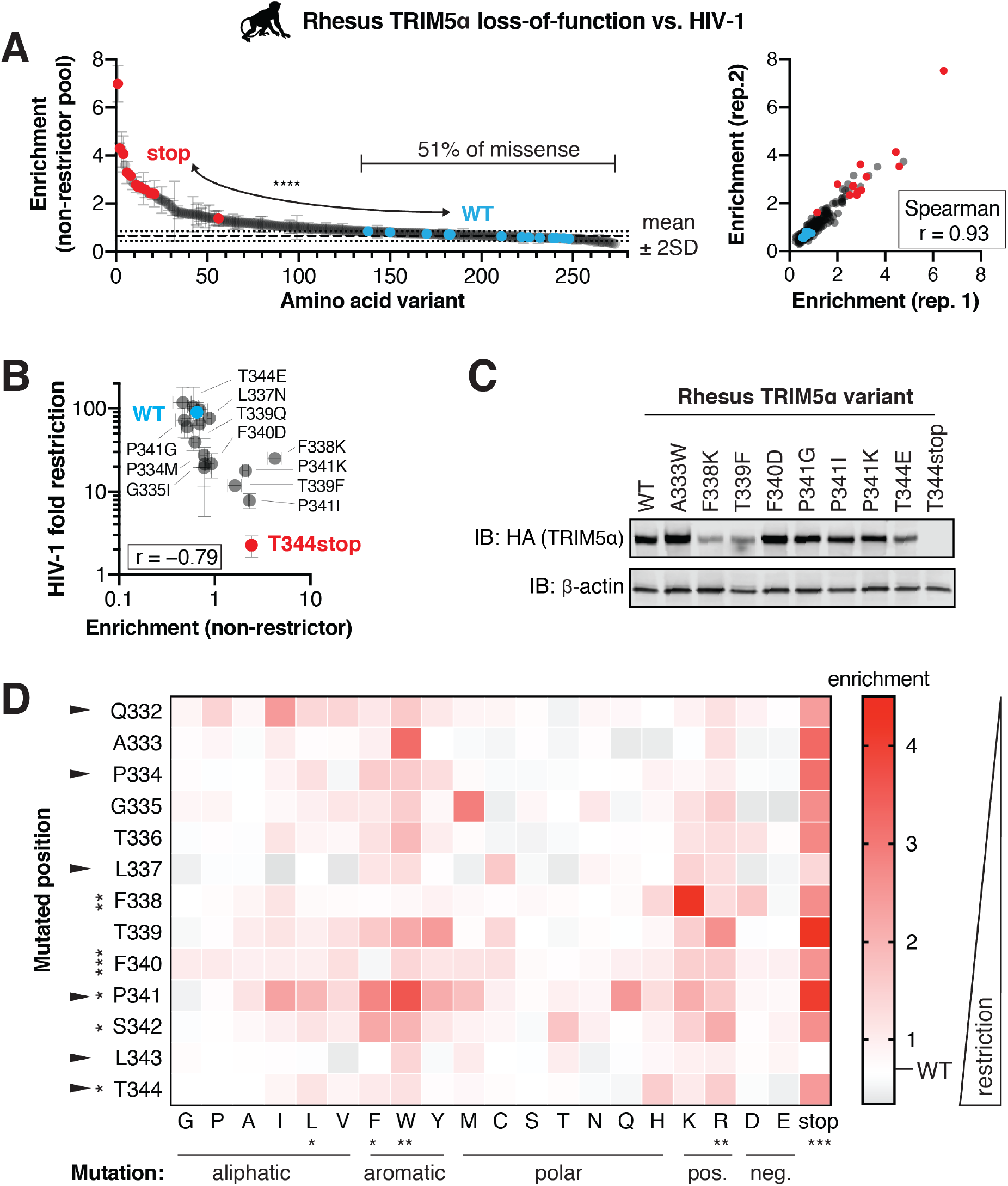
Rhesus macaque TRIM5α restriction of HIV-1 tolerates many mutations. A rhesus TRIM5α v1 DMS library was infected with HIV-1-GFP VLPs, and GFP-positive (non-restrictor) cells were sorted and sequenced. (**A**) Nonsense variants are highly enriched in the non-restrictor pool compared to WT (****p < 0.0001, student’s unpaired t-test with Welch’s correction), while half of all missense variants fall less than 2 SD above WT. Enrichment scores between two biological replicates are highly correlated. (**B**) Re-testing individual variants confirms that enriched variants have partially lost HIV-1 restriction, while depleted variants do not differ from WT. Spearman r; error bars, SD. (**C**) Steady-state levels of TRIM5α variants stably expressed in CRFK cells. (**D**) Enrichment in the HIV-1 non-restrictor pool relative to WT (white) for each variant, arrayed by position and amino acid mutation, is indicated by color intensity. The color scale was slightly compressed to avoid exaggerating a single mutant (L343stop) with enrichment > 4.5. Rapidly evolving sites are indicated with arrows. Statistical tests compare each position (across all variants) or each amino acid (across all positions) to WT, one-way ANOVA with Geisser-Greenhouse non-sphericity and Holm-Sidak’s multiple comparisons corrections; *p < 0.05, **p < 0.01, ***p < 0.001, ****p < 0.0001.

By re-testing individual variants, we confirmed that enrichment scores negatively correlate with antiviral potency (Figure 5B). We tested seven variants enriched for loss-of-restriction (more than 2 standard deviations above WT) and found that six lost HIV-1 restriction. The seventh variant (L337N) was only slightly outside the two standard-deviation cut-off for enrichment and correspondingly only slightly worse than WT in terms of HIV-1 inhibition. Thus, enrichment in the non-restrictor pool represents *bona fide* loss of restriction. All the rhesus TRIM5α variants we report here represent novel loss-of-function mutations. Their loss of HIV-1 restriction cannot be explained by loss of expression or protein stability (Figure 5C). For example, the F340D, P341I, and P341G variants were all expressed at WT levels but lost HIV-1 restriction. Moreover, the T344E variant retained restriction despite reduced expression levels.

We also re-tested ten rhesus TRIM5α variants not significantly enriched for loss-of-restriction (Figure 5B). Two variants (P334M, G335I) enriched one standard deviation above WT correspondingly retained only partial HIV-1 restriction relative to WT rhesus TRIM5α. The eight remaining variants retained HIV-1 inhibition, consistent with their lack of enrichment relative to WT. Based on this validation, we conclude that roughly half of all v1 loop single point mutations do not significantly reduce HIV-1 restriction by rhesus TRIM5α. Thus, a natural rhesus TRIM5α antiviral variant, much like the human TRIM5α-R332P variant, displays considerable mutational resilience.

We expected that conserved residues should be less tolerant of changes than rapidly evolving sites. However, we found that mutations in only 3 of 7 conserved sites, versus 2 of 6 rapidly evolving sites, led to significant loss of function (Figure 5D). Collectively, these results indicate that rhesus TRIM5α restriction of HIV-1 is highly robust to changes within the critical v1 loop at both rapidly evolving and conserved sites. The biochemical preferences for HIV-1 restriction are similar but not identical between rhesus and human TRIM5α. In both cases, the introduction of positive charge, particularly R, weakened HIV-1 inhibition (Figure 5D, compare to Figure 3C). In contrast, the introduction of bulky hydrophobic residues, including leucine (L), phenylalanine (F), and tryptophan (W), significantly impaired HIV-1 restriction by rhesus TRIM5α but did not affect the potency of human TRIM5α. These data suggest that both universal as well as lineage-specific requirements for the v1 loop shape TRIM5α restriction of HIV-1.

Our findings with TRIM5α restriction of HIV-1 suggest that single mutations can readily achieve gain-of-function. In contrast, loss-of-function mutations are not so abundant as to make adaptation unlikely. Thus, the evolutionary landscape of TRIM5α appears to resemble ‘rolling hills’ (Figure 1D) rather than rare, sharp peaks (Figure 1C).

### Resilience of antiviral restriction is a general property of TRIM5α adaptation

Our DMS analyses of human TRIM5α revealed unexpected ease of gaining antiviral potency against HIV-1 and potentially other lentiviruses. However, gains in potency against one virus might be offset by a concomitant loss of function against other viruses, as previously seen for the antiviral protein MxA (Colón-Thillet et al., 2019). Such functional tradeoffs might partially explain the evolutionary constraints acting on primate TRIM5α sequences. To explore this possibility, we investigated the mutational resilience of N-tropic murine leukemia virus (N-MLV) restriction by TRIM5α. N-MLV is strongly inhibited by both rhesus and human TRIM5α, and this activity is at least partly dependent on the v1 loop (Ohkura et al., 2006; Perron et al., 2006). We infected cells expressing either the rhesus (Figure 6A) or WT human TRIM5α (Figure 6B) v1 DMS libraries with GFP-marked N-MLV, sorted GFP-positive cells, and sequenced the non-restrictor variants. For both selections, stop codon variants were significantly more enriched than WT variants in the non-restrictor pool.

**Figure 6.**
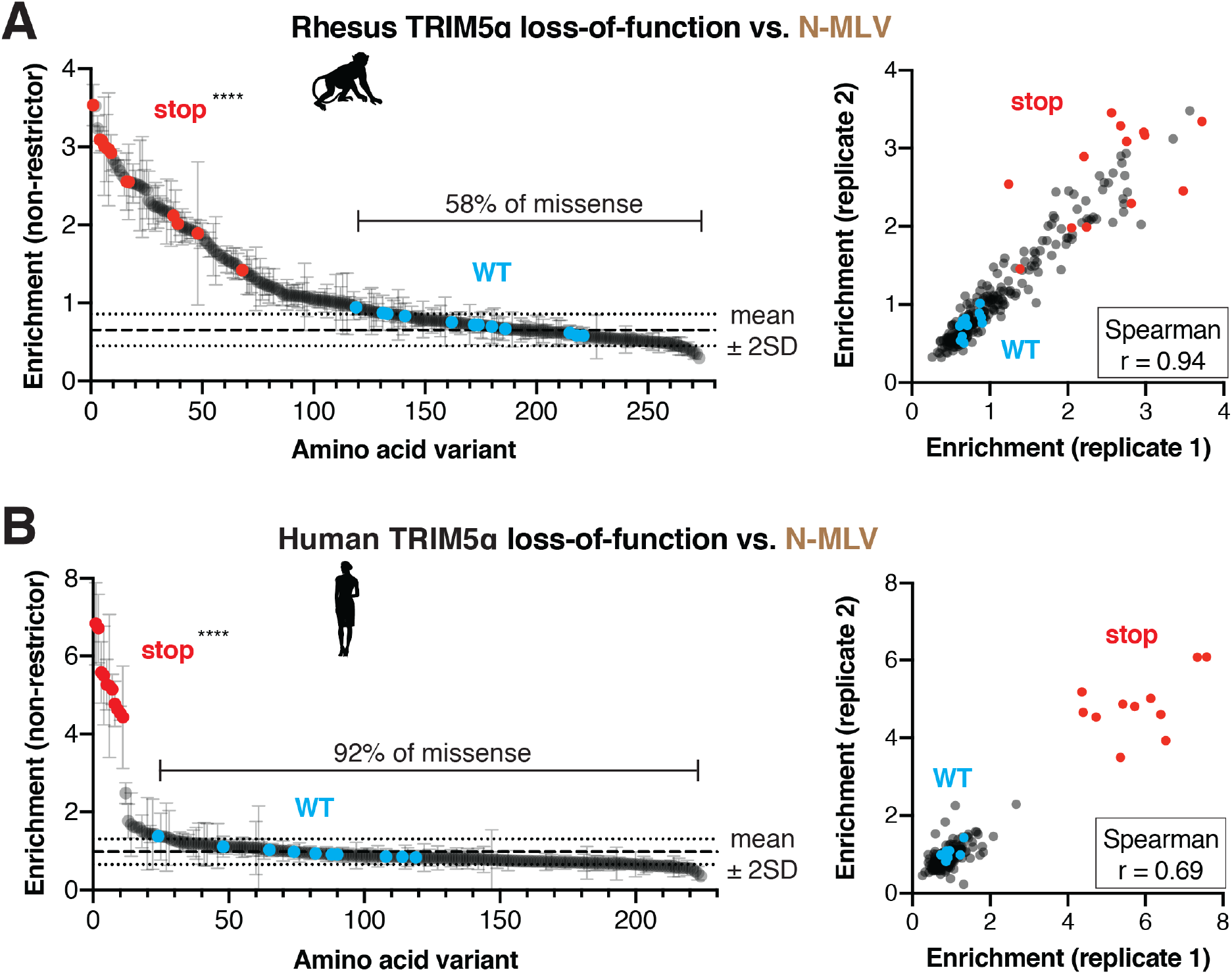
N-MLV restriction by primate TRIM5α is robust to single point mutants. The rhesus (**A**) or WT human (**B**) v1 DMS libraries were infected with N-MLV-GFP VLPs, and GFP-positive (non-restrictor) cells were sorted and sequenced. Nonsense variants are highly enriched in the non-restrictor pool compared to WT in both screens; ****p < 0.0001, student’s unpaired t-test with Welch’s correction. Error bars, SD. The fraction of all missense variants less than 2SD above WT mean is indicated. Enrichment scores between two biological replicates are highly correlated.

**Figure 6—figure supplement 1.**
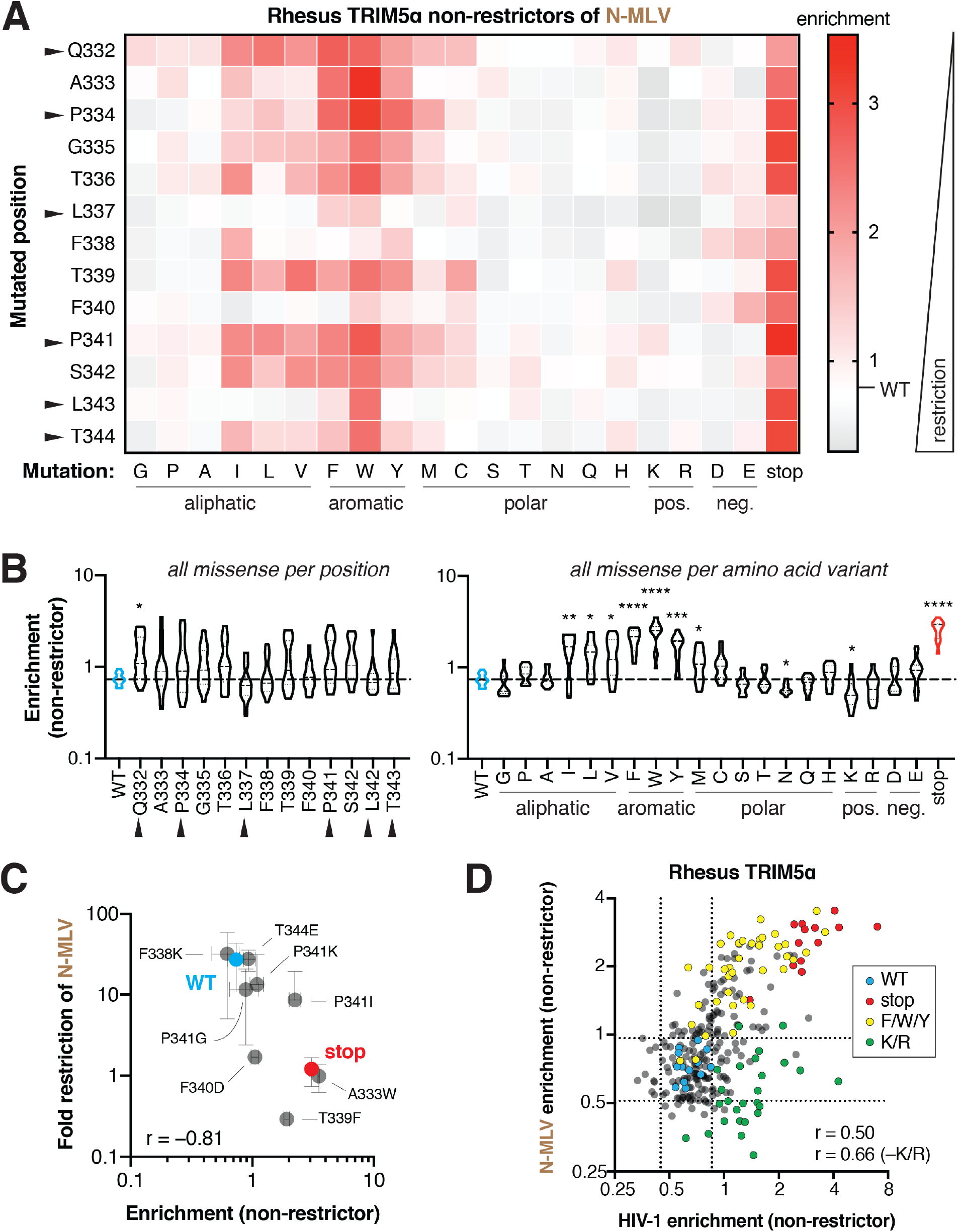
Biochemical preferences for rhesus TRIM5α restriction of N-MLV are distinct from HIV-1. (**A**) Enrichment scores in the N-MLV non-restrictor pool for each variant are arrayed by position and amino acid mutation. Enrichment (decreased restriction, red) relative to WT (white) is indicated by color intensity. (**B**) Missense variants at each position (across all variants) or each amino acid (across all positions); one-way ANOVA compared to WT with Geisser-Greenhouse non-sphericity and Holm-Sidak’s multiple comparisons corrections; *p < 0.05, **p < 0.01, ***p < 0.001, ****p < 0.0001. (**C**) Re-testing individual mutations confirms that enriched mutants have lost restriction (Spearman r = −0.81). Error bars, SD. (**D**) Rhesus TRIM5α restriction of HIV-1 and N-MLV has partially overlapping biochemical requirements. Positive charge (K/R, green) breaks only HIV-1 restriction, whereas stop codons (red) and aromatic residues (F/W/Y, yellow) weaken restriction of both viruses. Excluding K and R improves the correlation compared to all variants (Spearman r = 0.66 or 0.50, respectively). Dashed lines indicate 2 SD above and below the mean enrichment for WT variants (blue) in each screen.

Similar to HIV-1 restriction, we found that most missense mutations (143, 58%) in rhesus TRIM5α were tolerated for N-MLV restriction (Figure 6A). However, some missense mutations dramatically reduced N-MLV restriction, affirming that the v1 loop is indeed critical for inhibition of N-MLV (Figure 6—figure supplement 1A-C). In particular, hydrophobic and especially aromatic residues at most positions in the v1 loop significantly decreased N-MLV restriction. This preference against aromatic residues is similar between HIV-1 and N-MLV restriction. However, N-MLV restriction is insensitive to the introduction of positively charged residues, which disrupt HIV-1 inhibition (Figure 6—figure supplement 1D). These results indicate that the evolutionary landscape for rhesus TRIM5α against N-MLV is distinct from that of HIV-1. Nevertheless, the overall degree of mutational resilience against both viruses is remarkably similar: less than half of all missense mutations disrupt restriction of either virus.

Human TRIM5α restriction of N-MLV was even more resilient to mutation than rhesus TRIM5α. Almost all variants (187, 92%) had no effect on N-MLV restriction (Figure 6B, Figure 6–figure supplement 2). Indeed, our selection for non-restrictive variants only strongly enriched for stop codons. This extreme mutational resilience may reflect the massive potency (>250-fold restriction, data not shown) of human TRIM5α against N-MLV, and/or a decreased reliance on the v1 loop for N-MLV recognition by human TRIM5α (Perron et al., 2006). We validated several human TRIM5α mutants as retaining nearly WT levels of N-MLV restriction (Figure 6—figure supplement 2C). Thus, both rhesus and human TRIM5α inhibition of N-MLV is highly resistant to mutations, allowing mutational flexibility without loss of pre-existing antiviral restriction. These results, in conjunction with the substitution tolerance of HIV-1 restriction by human R332P and WT rhesus TRIM5α, indicate that mutational resilience is a general property of TRIM5α’s rapidly evolving v1 loop.

**Figure 6—figure supplement 2.**
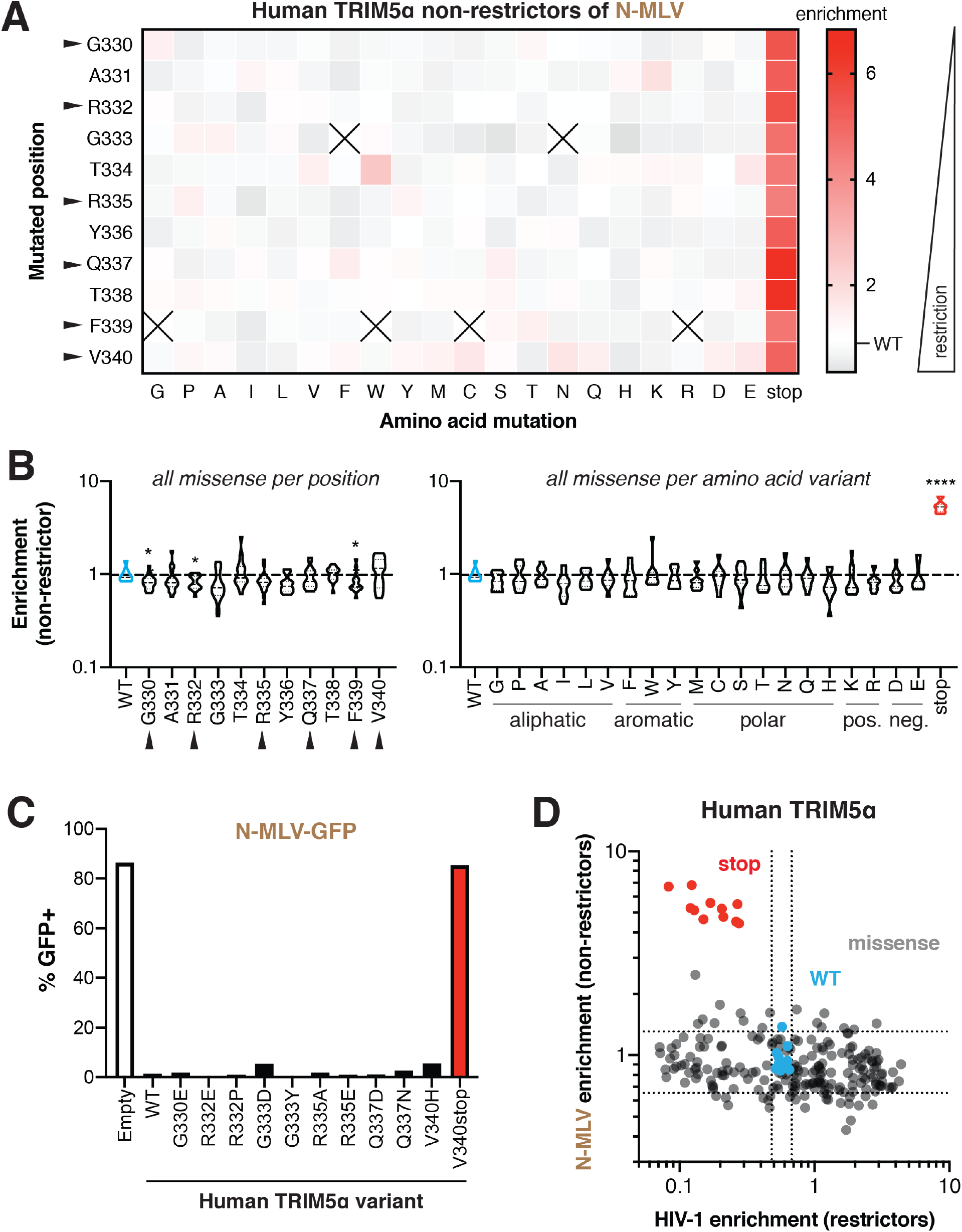
Missense mutations do not disrupt human TRIM5α restriction of N-MLV. (**A**) Enrichment scores in the N-MLV non-restrictor pool for each variant are arrayed by position and amino acid mutant. Enrichment (decreased restriction, red) relative to WT (white) is indicated by color intensity. Variants marked with X were excluded due to low input representation. Rapidly evolving residues are indicated with black arrows. (**B**) Missense variants at each position (across all variants) or each amino acid (across all positions); one-way ANOVA compared to WT with Geisser-Greenhouse non-sphericity and Holm-Sidak’s multiple comparisons corrections; *p < 0.05, **p < 0.01, ***p < 0.001, ****p < 0.0001. (**C**) Re-testing individual mutations confirms that mutations have little effect on N-MLV restriction. Results are representative of at least 3 independent experiments. (**D**) HIV-1 gain-of-restriction compared to N-MLV loss-of-restriction. No anti-correlation is evident, with the exception of stop codons (red). Dashed lines indicate 2 SD above and below the mean enrichment for WT variants (blue) in each screen.

## DISCUSSION

Antiviral restriction factors are locked in high-stakes tit-for-tat evolutionary arms races with target viruses. However, viruses would appear to have the upper hand in these battles because of their higher mutation rates, shorter generation times, and larger population sizes. Although host genomes have the advantage of encoding a diverse, polygenic immune response, evolutionary constraints acting on innate immune genes could curtail their adaptive potential. Here, using deep-mutational scanning approaches combined with viral infection assays, we investigated the evolutionary landscape of adaptation of the most rapidly evolving segment, the disordered v1 loop, of the retroviral restriction factor TRIM5α. We focused on this loop because of its critical role in adapting to changing viral repertoires.

We found two attributes of this evolutionary landscape that favor host immune evolution. First, human TRIM5α readily gains significant HIV-1 restriction: roughly half of all single missense mutations allow human TRIM5α to better restrict HIV-1 (Figures 2-3). Based on our results, we infer that positive charge is the dominant impediment to HIV-1 inhibition in human TRIM5α (Figure 3D). Removal of this positive charge improved human TRIM5α restriction not only of HIV-1 but also of multiple lentiviruses (Figure 4A). Recent findings revealed that cyclophilin A (CypA) protects the HIV-1 capsid from TRIM5α recognition (Kim et al., 2019; Selyutina et al., 2020; Veillette et al., 2013). Although structural studies currently lack sufficient resolution to observe the molecular details of the TRIM5α-capsid interaction, we speculate that positive charge in the v1 loop impairs an interaction between TRIM5α and the capsid’s CypA-binding site via electrostatic repulsion. Given the almost universally detrimental impact of positive charge on lentiviral restriction, the fixation of positive charge during the evolution of hominid TRIM5α is perplexing (Figure 1A). We have not identified a candidate virus driving this fixation, for which we might expect a positive charge in the v1 loop to be beneficial. However, since the critical R332 was fixed at the common ancestor of humans, chimps, and bonobos ~10 million years ago, it is possible that this viral challenge was a paleovirus that has since gone extinct. Another possibility is that positive charge did not aid in gaining new antiviral specificity, but rather has a function independent of viral capsid recognition, such as suppressing innate immune signaling in the absence of infection (Lascano, Uchil, Mothes, & Luban, 2015; Pertel et al., 2011; Tareen & Emerman, 2011).

Surprisingly, our comprehensive DMS analyses also revealed that both rapidly evolving and conserved residues can contribute to antiviral adaptation (Figure 3B). For instance, many variants at position 333 of human TRIM5α led to increased restriction of HIV-1 and other lentiviruses (Figures 3A, 4A). Thus, it is unclear why simian primates have retained a glycine at this position (333 in human, 335 in macaques). One possibility is that changes in this residue might be generally deleterious for TRIM5α function, yet subsequent analyses revealed little or no impairment of TRIM5α antiviral functions (Figure 5D, Figure 6—figure supplement 1A, Figure 6—figure supplement 2A). This suggests that conservation of G333 either reflects some cellular constraint or was recurrently selected for by a viral lineage distinct from the viruses we tested here (HIV-1 and N-MLV).

Our analyses uncovered a second unexpected, advantageous aspect of TRIM5α’s evolutionary landscape: its antiviral restriction displays remarkable mutational resilience across multiple orthologs and against two divergent retroviruses. 51-92% of all possible missense variants retain antiviral activity (Figure 6—figure supplement 3). This resilience is manifest even when potent antiviral activity is newly acquired via a single mutation, as with the R332P variant of human TRIM5α against HIV-1. Therefore, we conclude that the fitness landscape of TRIM5α’s rapidly evolving v1 loop resembles ‘rolling hills’ (Figure 1D), in which valleys are infrequent and only one evolutionary step removed from mutationally-tolerant plateaus.

TRIM5α’s permissive landscape contrasts with the relative inflexibility of ligand-binding domains in the core of evolutionarily conserved proteins (Guo et al., 2004; McLaughlin et al., 2012; Suckow et al., 1996). However, these studies found increased mutational tolerance in peripheral, disordered loops, such as those employed for viral ligand binding by TRIM5α as well as MxA (Mitchell et al., 2012). The use of flexible loops thus grants rapidly evolving restriction factors mutational flexibility without significant risk of disrupting core protein structure. Intriguingly, TRIM5α’s reliance on the v1 loop for specificity mirrors that of antibodies’ dependence on complementarity-defining loops for antigen recognition. Indeed, a high degree of mutational tolerance within complementarity-defining loops allows somatic hypermutation to significantly increase antibody-antigen affinity (P. S. Daugherty, Chen, Iverson, & Georgiou, 2000; Sheng et al., 2017).

**Figure 6—figure supplement 3.**
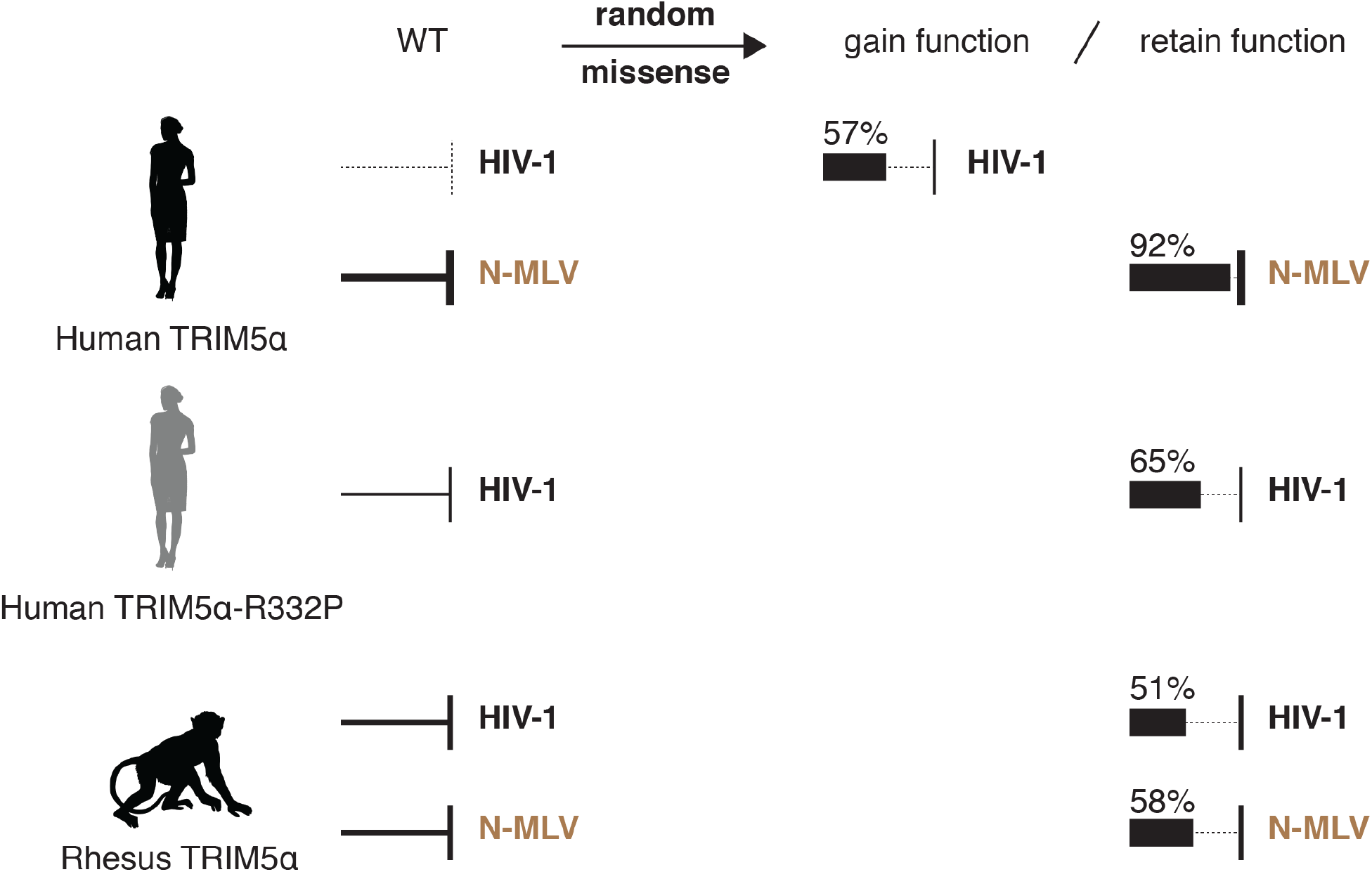
Summary of deep mutational scanning results. Random missense mutations in human TRIM5α frequently improve HIV-1 restriction. Multiple TRIM5α orthologs, including a *de novo* HIV-1 restrictor (human TRIM5α-R332P), display high mutational resilience against two distantly related retroviruses. Arrow thickness represents antiviral potency; the fraction of missense mutants that gain or retain function are indicated.

Overall, our analyses reveal not only many paths for TRIM5α to gain antiviral function but also an unexpectedly low probability of losing antiviral function via single mutations. Such landscapes should be highly advantageous to host genomes in evolutionary arms races with viruses. Mutational tolerance allows the accumulation of neutral variants that do not compromise antiviral function among antiviral genes in a population. Many of these novel variants may carry the capacity to restrict additional viruses, whether these result from cross-species transmissions or mutations that allow species-matched viruses to evade recognition by the dominant antiviral allele. Indeed, mutational tolerance has been shown to facilitate the evolution of *de novo* functions through the accumulation of neutral mutations (Draghi, Parsons, Wagner, & Plotkin, 2010; Hayden, Ferrada, & Wagner, 2011). Although rare in human populations (Clarke et al., 2017), extensive polymorphism within the v1 loop of TRIM5α in Old World monkeys results in diverse antiviral repertoires that have been maintained by balancing selection (Newman et al., 2006). Thus, antiviral proteins such as TRIM5α appear to evolve with low-cost, high-gain fitness landscapes that favor their success in co-evolutionary battles with rapidly evolving retroviruses.

## MATERIALS AND METHODS

### Plasmids and cloning

All virus-like particles (VLPs) were generated using three plasmids to ensure a single round of infectivity: a pseudotyping plasmid for transient expression of the VSV-G envelope protein (pMD2.G, Addgene plasmid #12259, gift from Didier Trono), a plasmid for transient expression of the viral gag/pol, and a transfer vector encoding a green fluorescent protein integration reporter between the corresponding viral LTRs. HIV-1 VLPs were made with the transfer vector pHIV-ZsGreen (B. E. Welm, Dijkgraaf, Bledau, Welm, & Werb, 2008); HIV-2 and SIV VLPs used the pALPS-eGFP transfer vector (McCauley et al., 2018); and N-MLV was made with pQCXIP-eGFP, encoding GFP between the EcoRI and ClaI sites of pQCXIP (P.S. Mitchell, unpublished). HIV-1 VLPs were made with p8.9NdSB bGH BlpI BstEII, encoding the NL4.3 HIV-1 gag/pol (Berthoux, Sebastian, Sokolskaja, & Luban, 2004). HIV-2 VLPs used a chimeric gag/pol, in which the HIV-1 CA sequence (residues 1-202) was replaced by HIV-2_ROD_ (p8.9NdSB bGH BlpI BstEII HIV-2 CA) (Pizzato et al., 2015). For SIV VLPs, pCRV1-based gag/pol chimeric vectors replaced the HIV-1 CA-NTD (residues 1-146) with the corresponding residues of either SIVmac239, a virus passaged in rhesus macaques, here SIVmac (pHIV-MAC, containing an A77V mutation); or SIVcpzGab2, a natural isolate from chimpanzee, here SIVcpz (pHIV-Gb2) (Kratovac et al., 2008). N-MLV VLPs were generated using pCIG3N, encoding the N-MLV gag/pol (Bock, Bishop, Towers, & Stoye, 2000). For stable expression of TRIM5α constructs from pQCXIP, the MLV gag/pol was transiently expressed from JK3; the pseudotyping envelope protein was transiently expressed from the L-VSV-G plasmid and driven by expression of Tat from the CMV-Tat plasmid.

C-terminally HA-tagged human, human with the rhesus macaque v1 loop, and rhesus macaque TRIM5α (Sawyer et al., 2005) were amplified and cloned into pQCXIP (Takara Bio, Kusatsu, Shiga, Japan), just upstream of the IRES–puromycin resistance cassette, between the EcoRI and NotI restriction sites. Targeted TRIM5α mutations were generated by Quikchange PCR using primers containing the desired point mutation flanked by 17-25 nucleotides of homology on each side of the mutation. Primestar polymerase (Takara Bio) was used to minimize errors during full-plasmid amplification, followed by DpnI digestion of unmodified parent DNA. All plasmids were cloned into high-efficiency chemically competent DH5α (NEB, Ipswich, MA, USA). Plasmids were purified using PureYield miniprep kits (Promega, Madison, WI, USA), and coding sequences were verified by complete sequencing. See Table 1 for all primers used in cloning and sequencing.

**Table 1.**
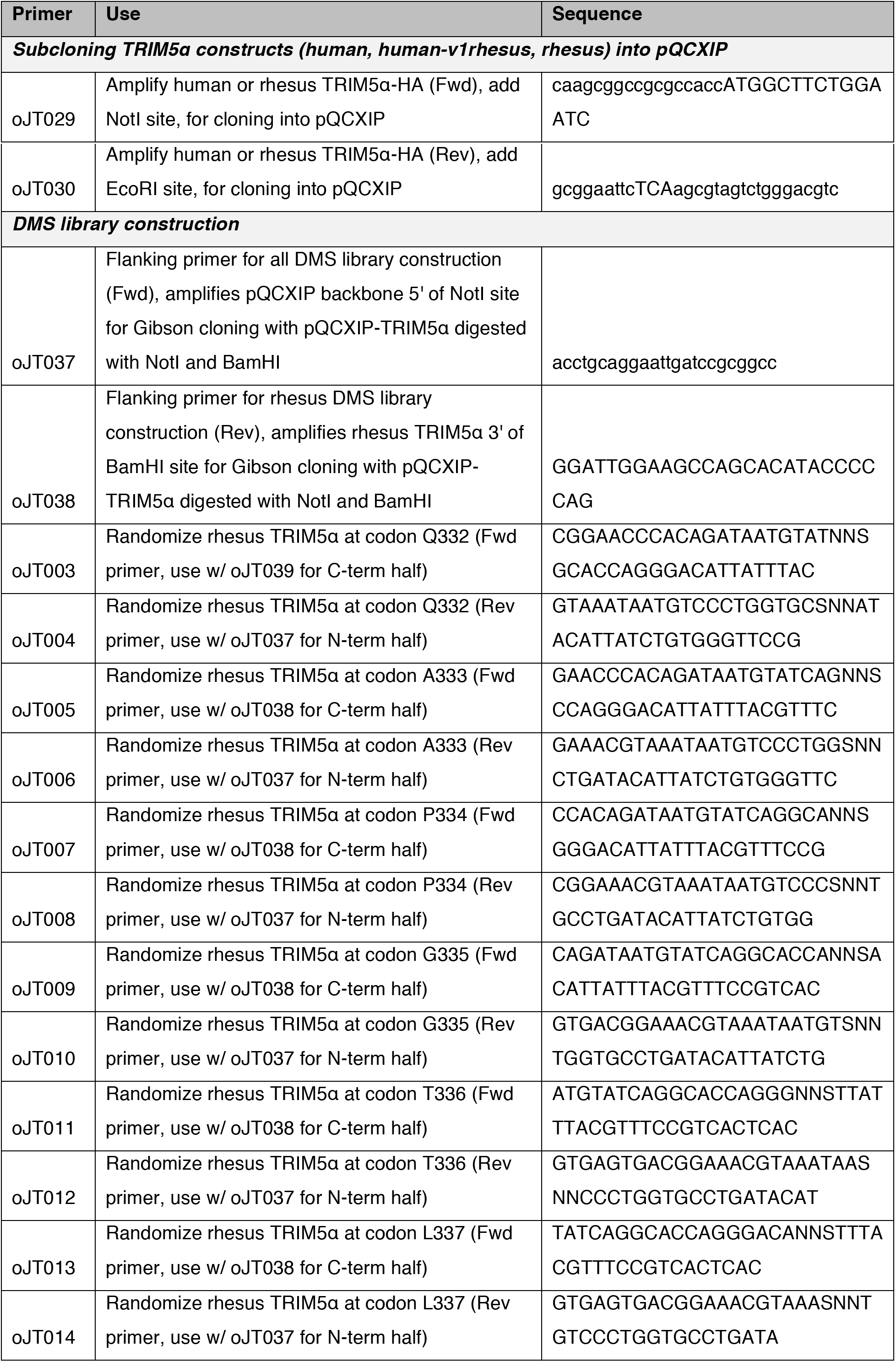
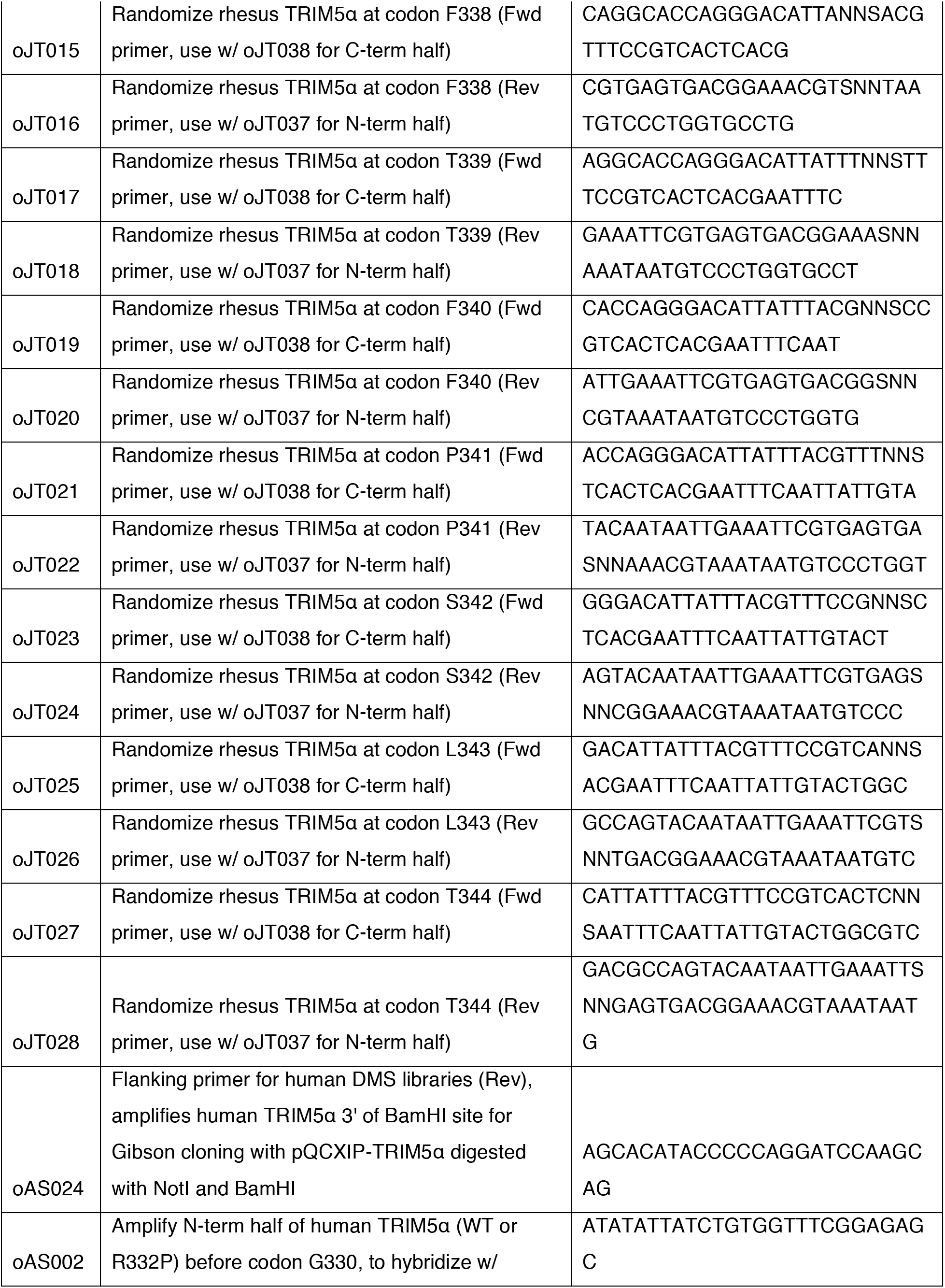
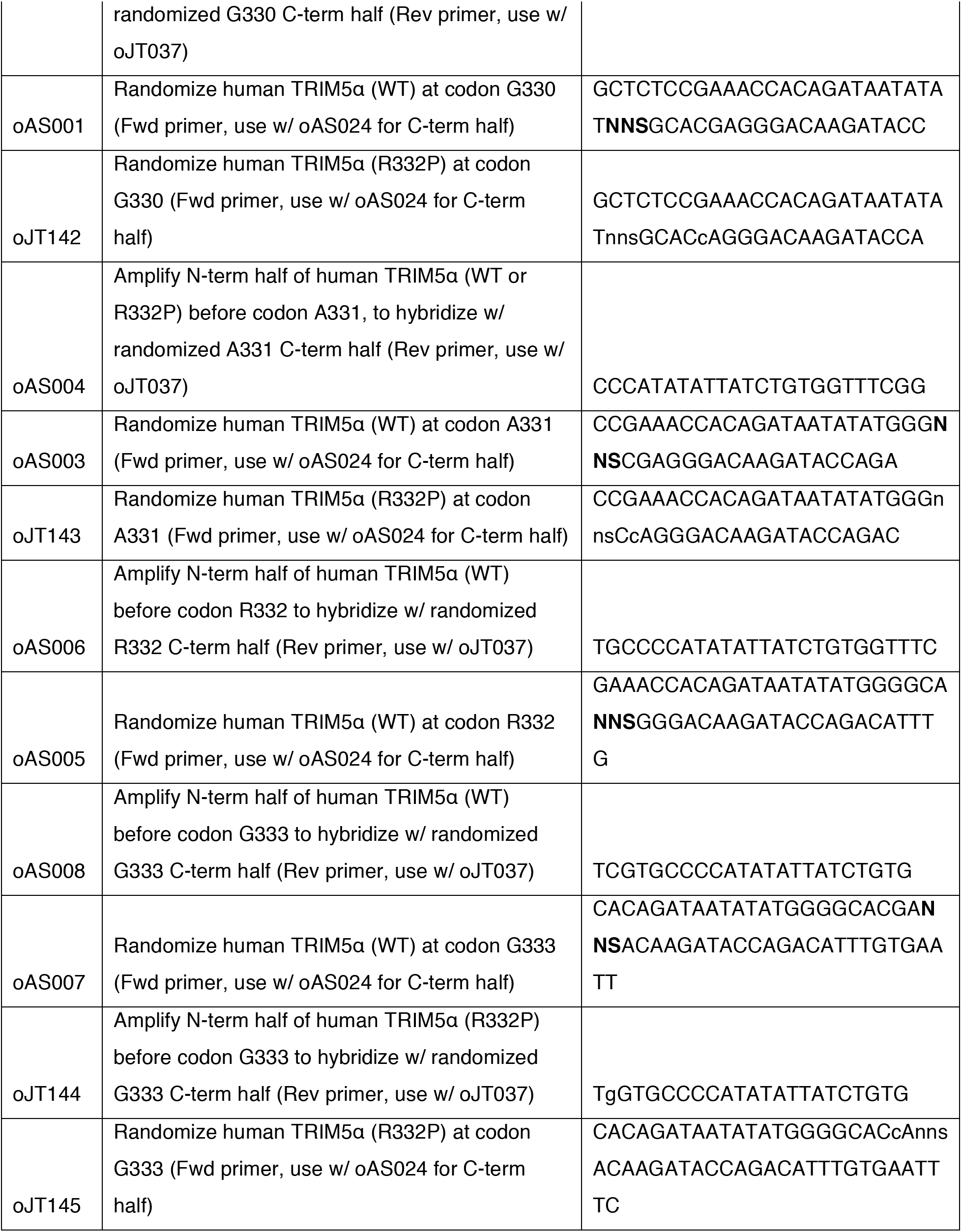
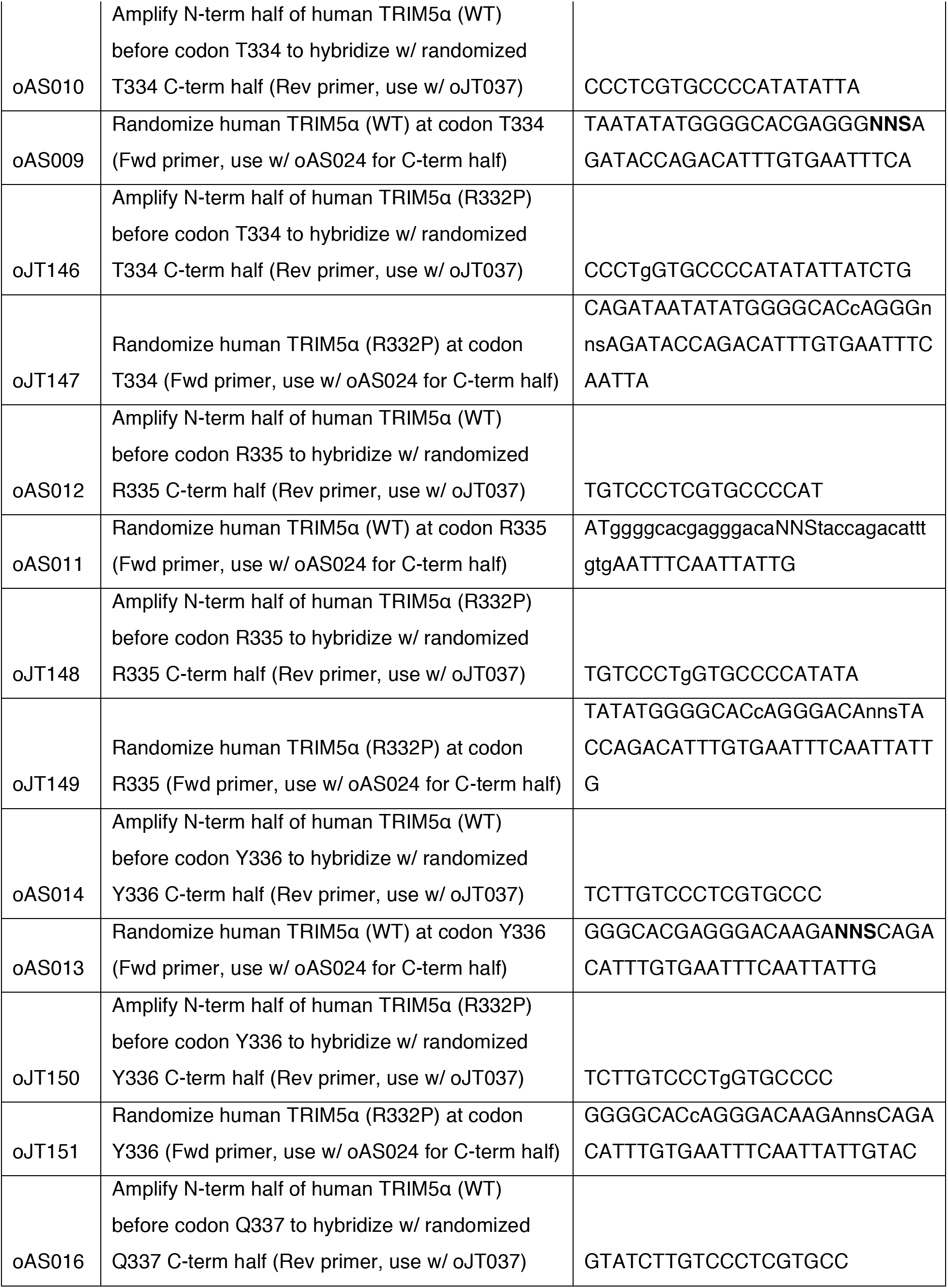
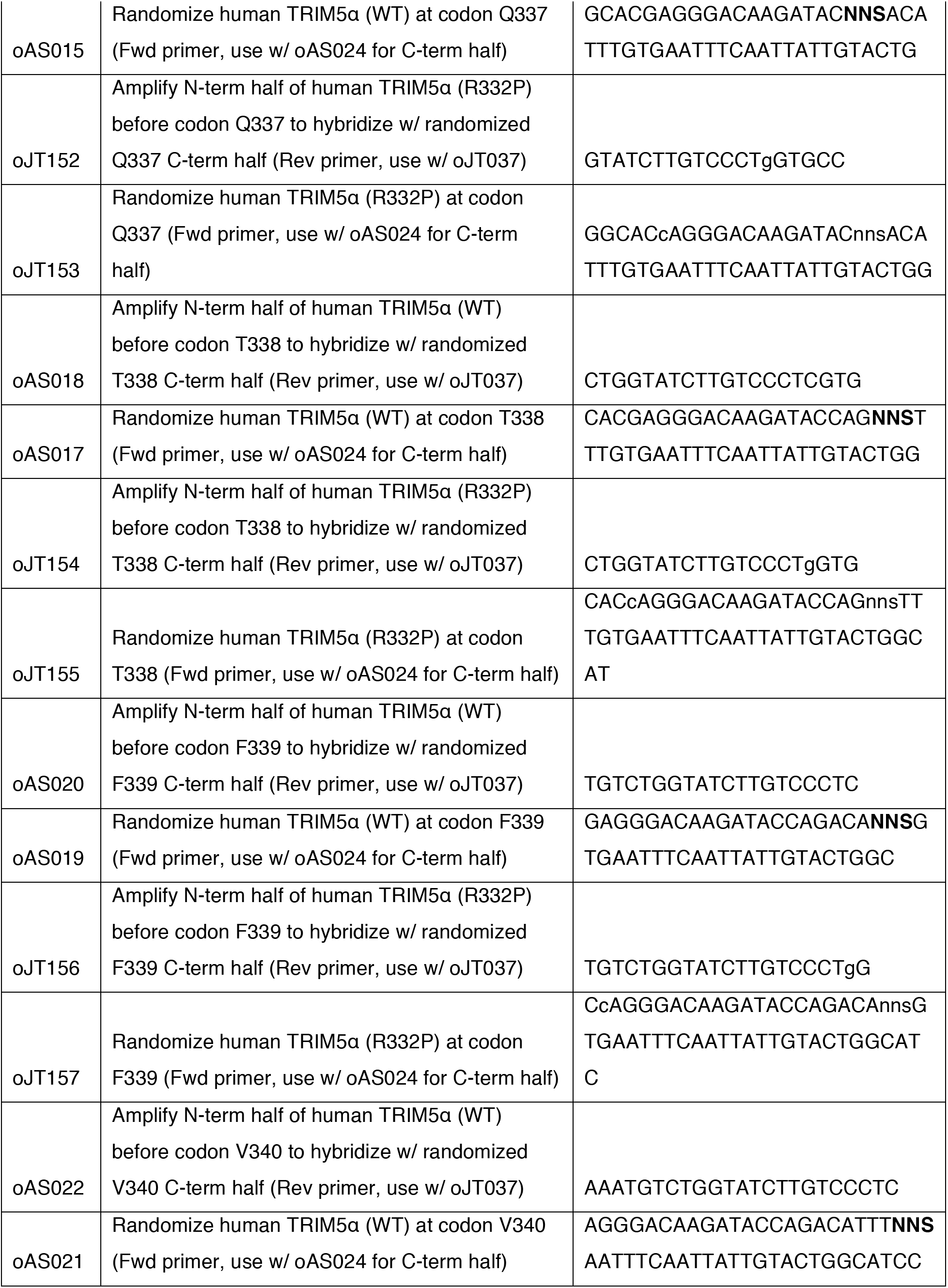
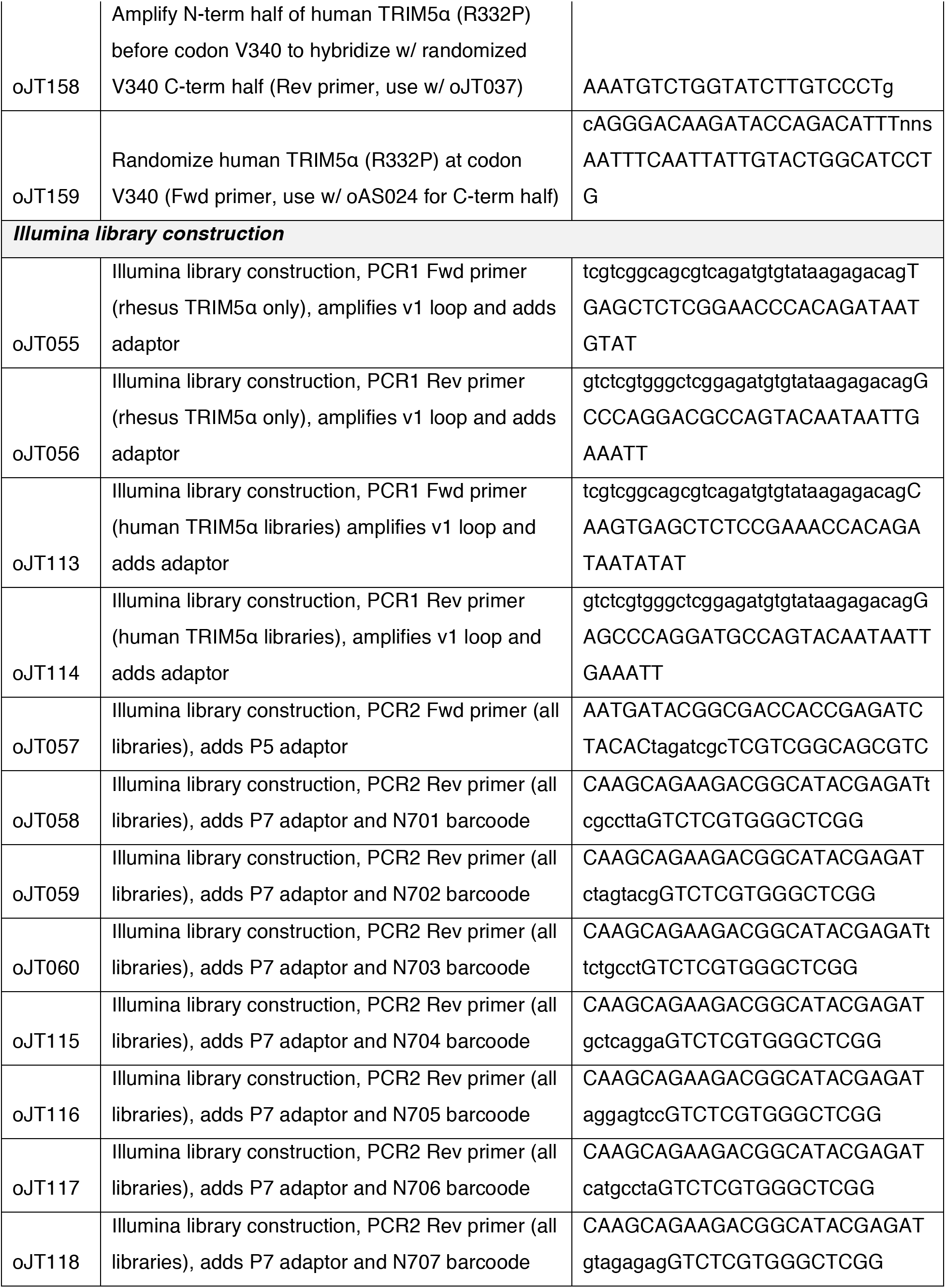
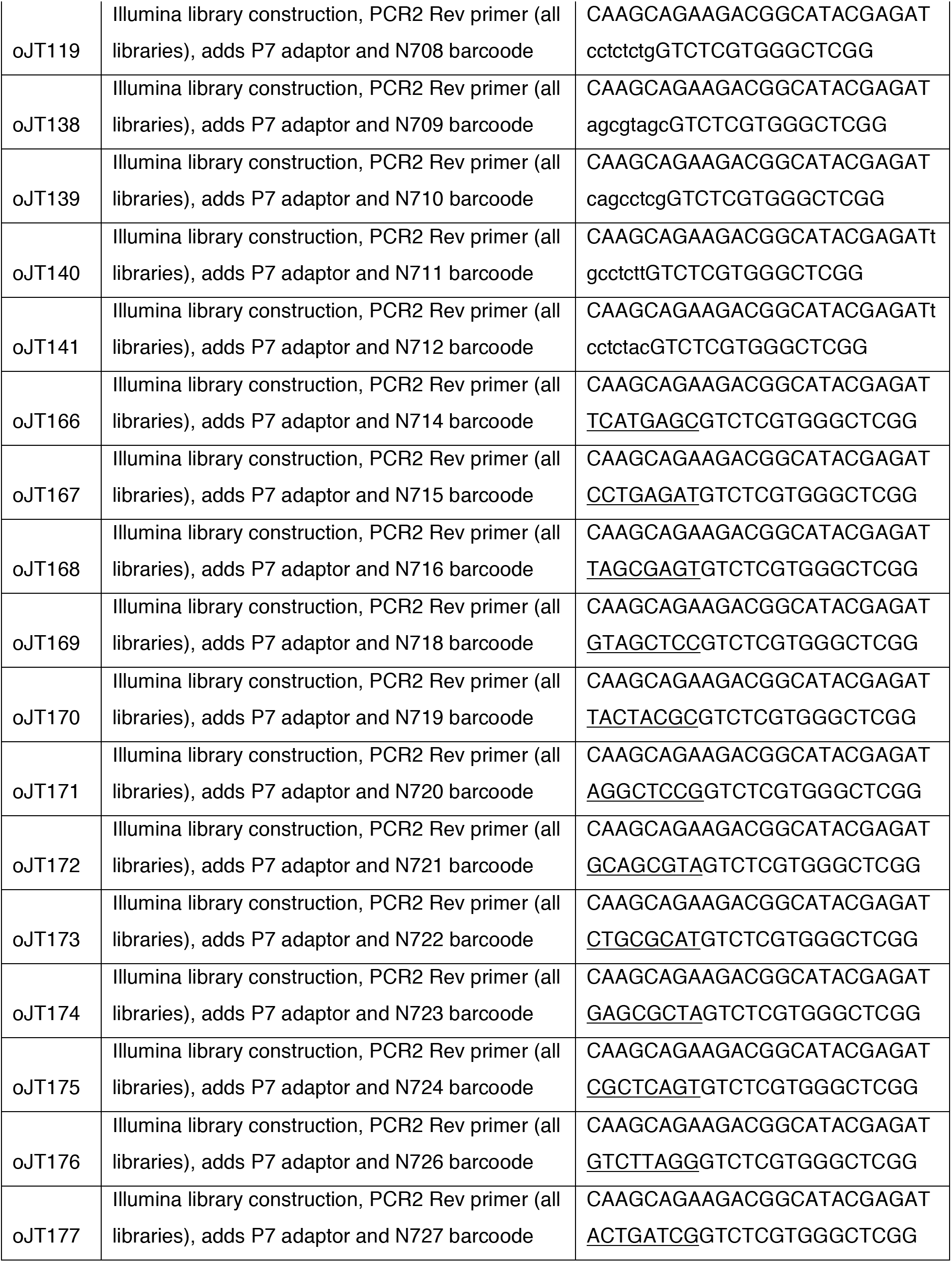
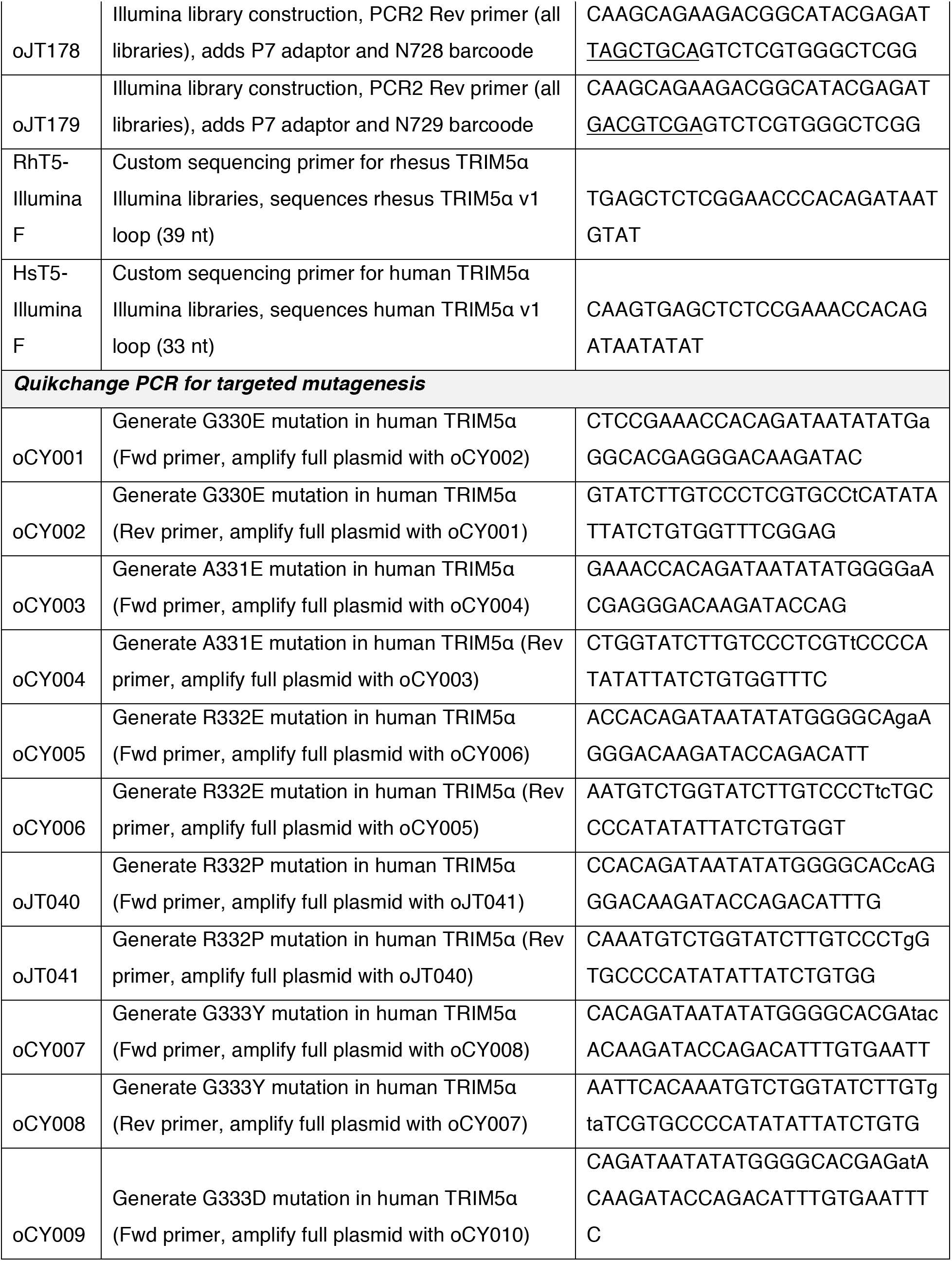
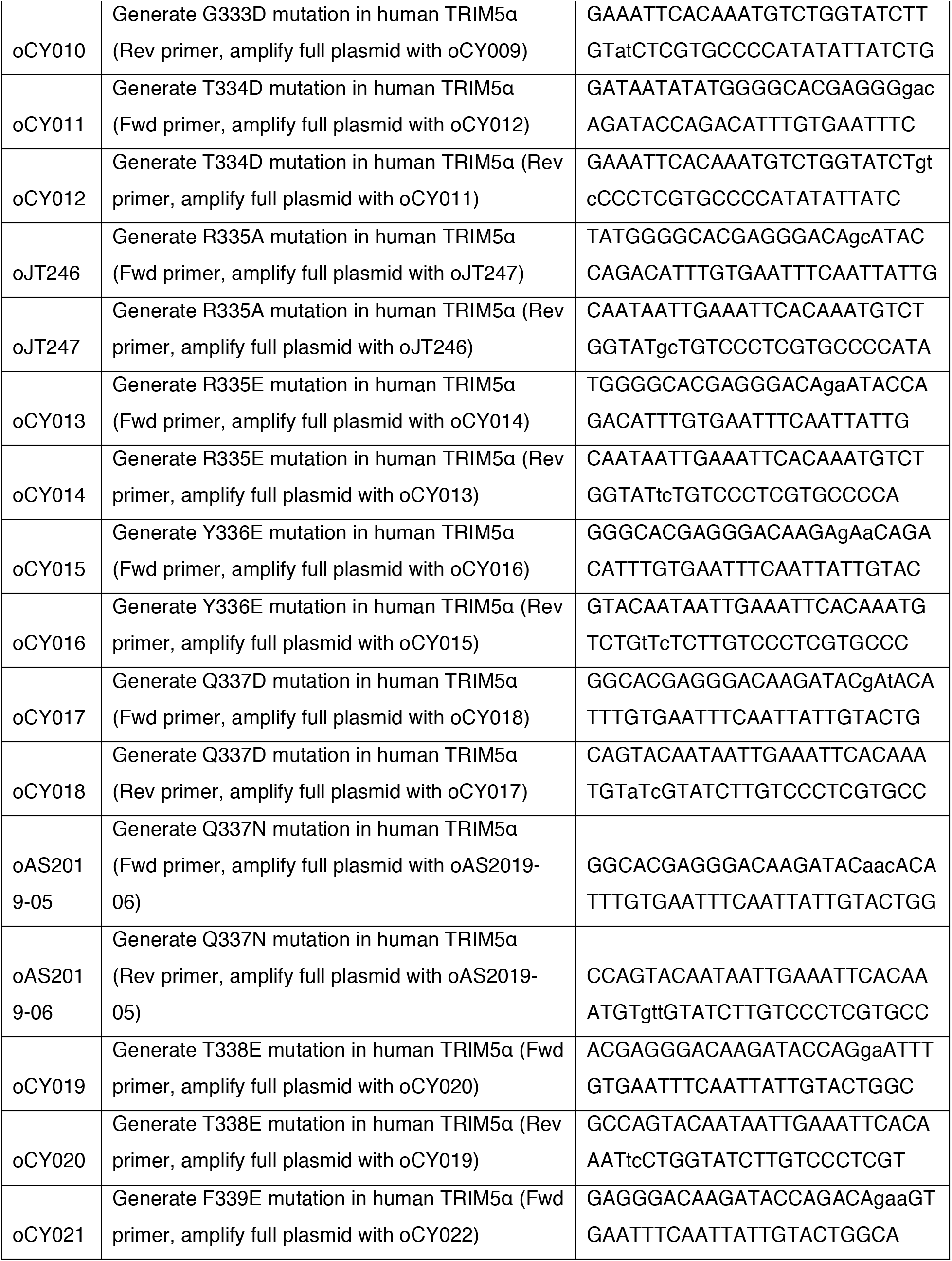
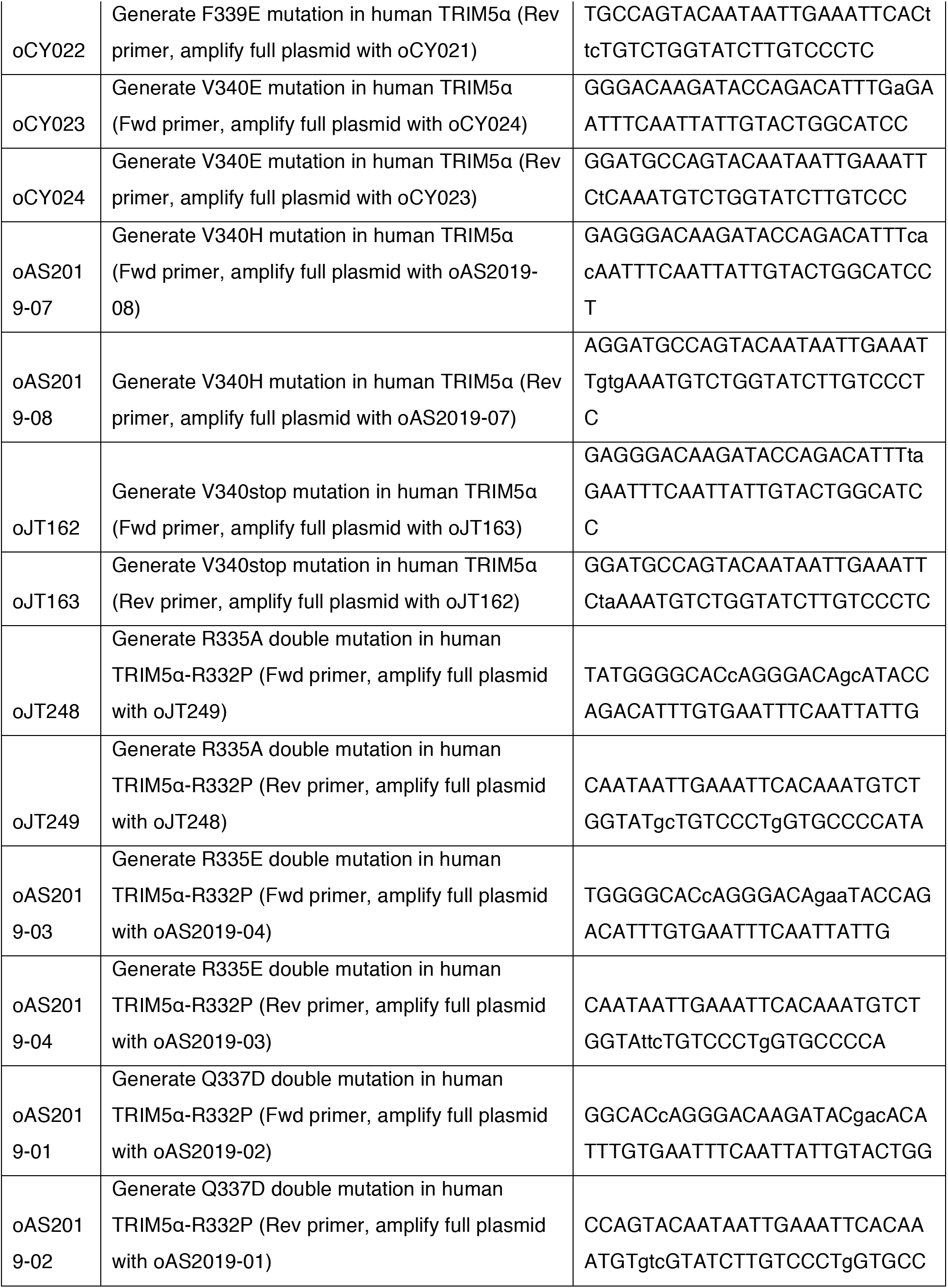
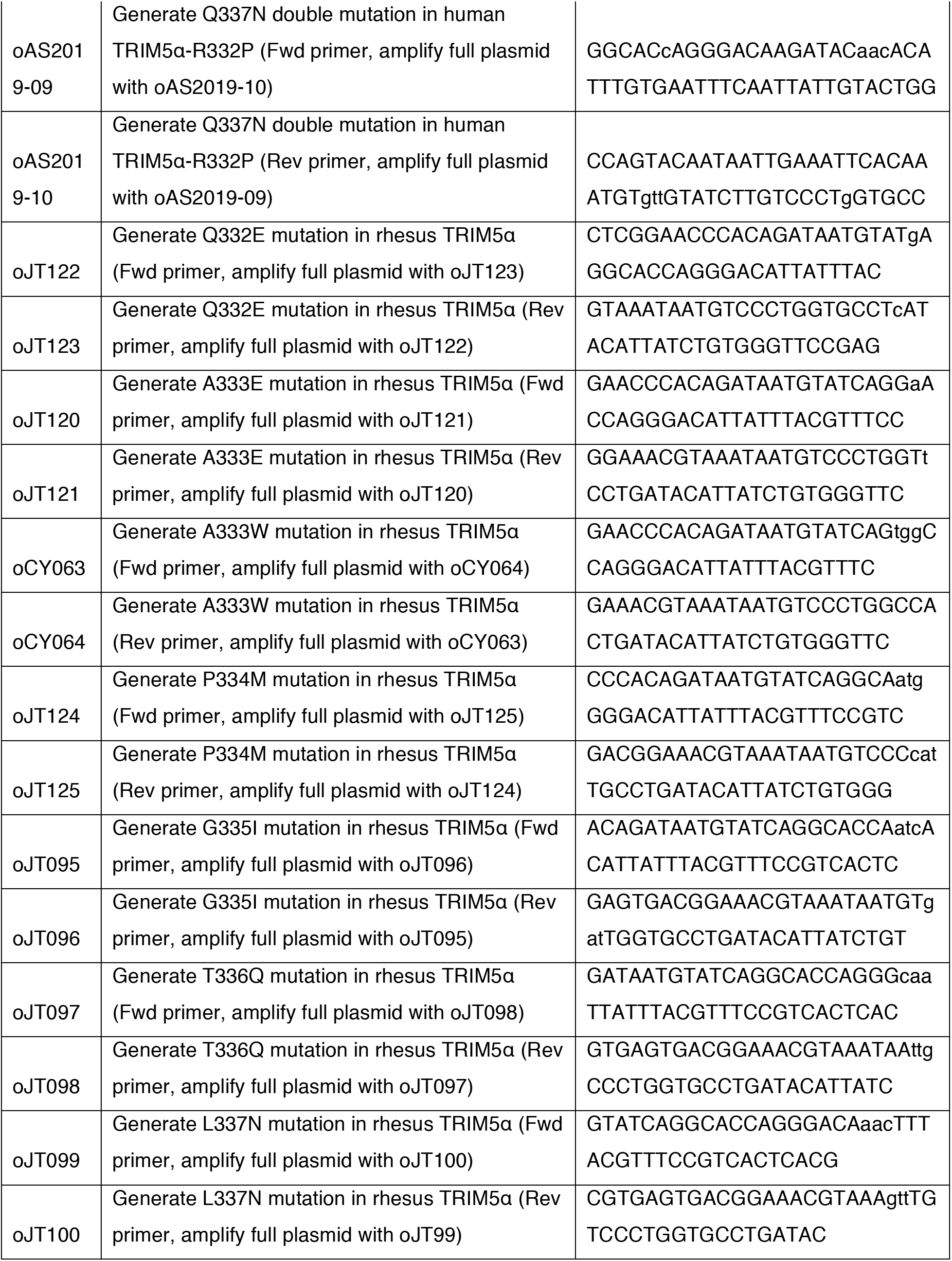
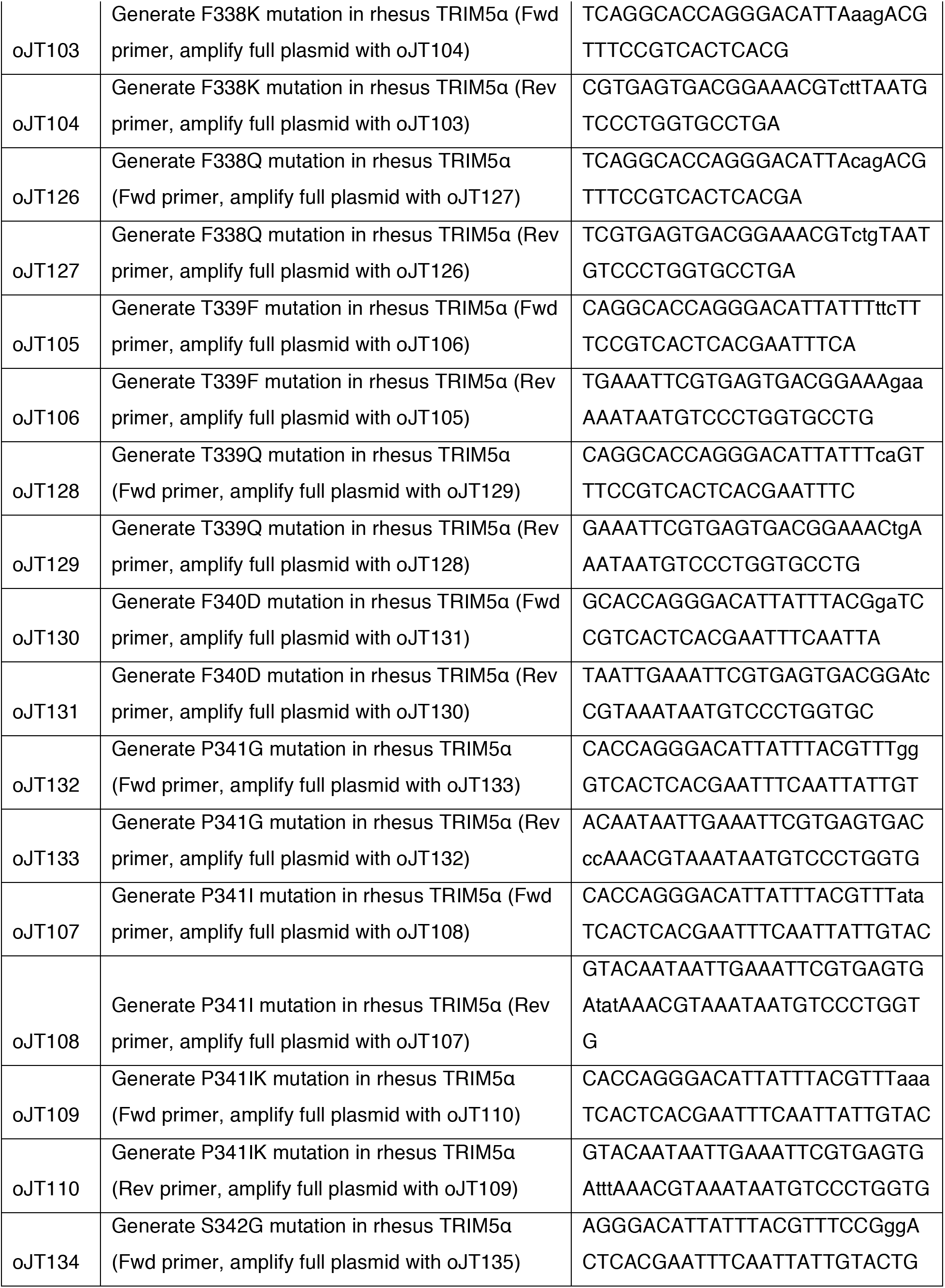
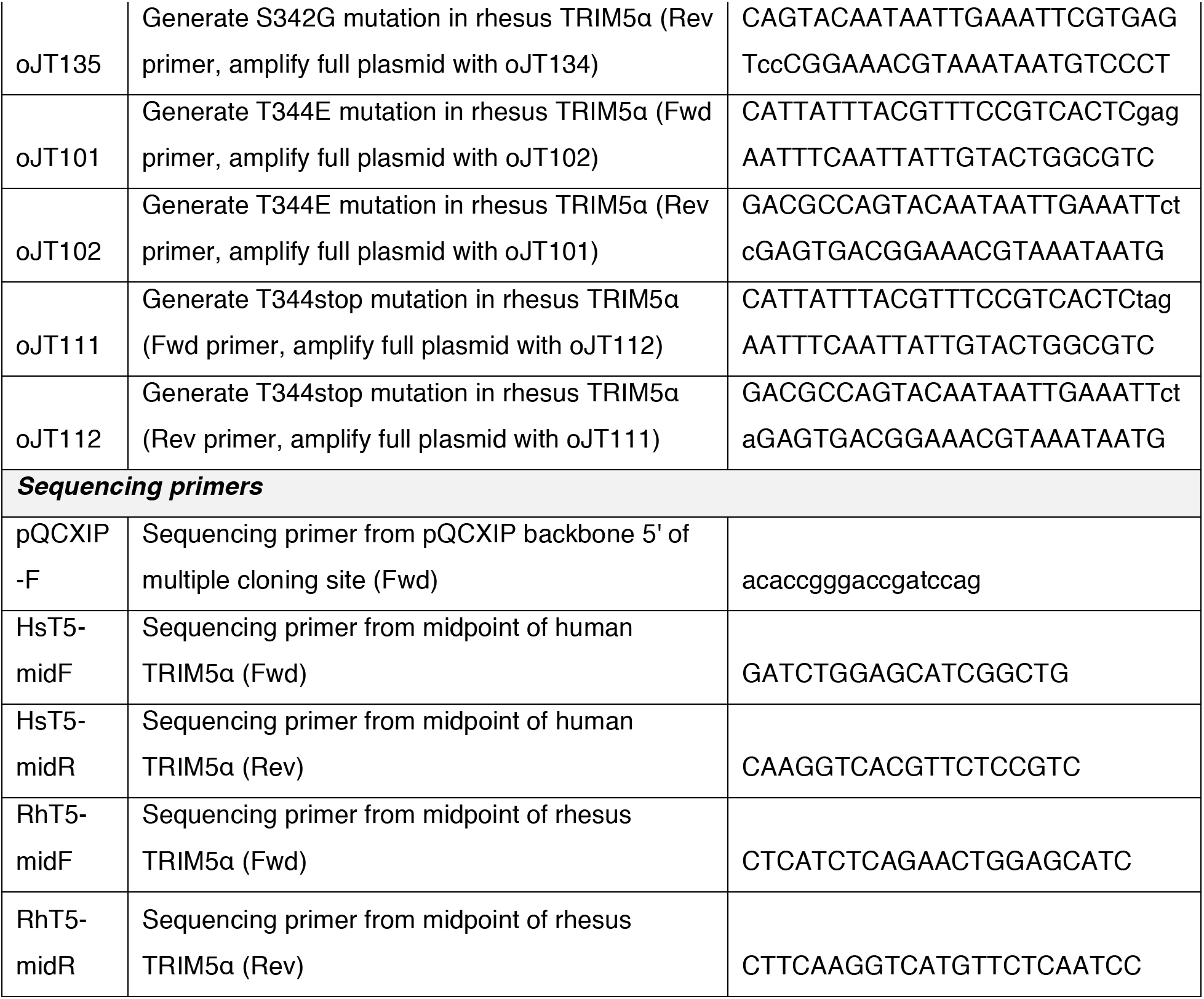
Primers used in this study.

Deep mutational scanning libraries were generated using degenerate primers to amplify TRIM5α-HA in pQCXIP using high-fidelity Q5 polymerase (NEB). Degenerate primers contained a single NNS codon (N = A/T/C/G, S = C/G), which encodes all 20 amino acids with only 1 stop codon among 32 possibilities. For each of the 11 or 13 codons in the v1 loop of human or rhesus TRIM5α, respectively, the two halves of TRIM5α were amplified separately with shared flanking primers and unique internal primers for each codon (Table 1). For the human TRIM5α library in which R332P was fixed, internal primers matched the R332P variant of TRIM5α and codon 332 was not randomized. Internal primers encoded NNS at the designated codon flanked by 17-25 nucleotides of homology on each side; the forward and reverse internal primers shared 17-25 nucleotides of homology to promote hybridization between the N- and C-terminal PCR fragments. The codon-matched N- and C-terminal fragments were combined and amplified into a single fragment using the same flanking primers as in the first amplification. PCR products were gel purified and cloned via Gibson assembly (NEB) into pQCXIP-TRIM5α-HA of the matching species, which had been digested with NotI and BamHI and gel purified. Gibson assembly products were transformed into high-efficiency chemically competent DH5α (NEB) with 30 minutes of heat shock recovery. Serial dilutions were plated to count the number of unique colonies, and transformations were repeated until at least 100x library coverage was achieved (human: 32 x 11 x 100 = 3.52 x 10^4^ colonies; rhesus: 32 x 13 x 100 = 4.16 x 10^4^ colonies). To ensure library quality, 40 random colonies were sequenced from each library. Clones were verified to have insert by analytical restriction digest, and the coding sequence was fully sequenced to ensure that (1) each clone had only 1 mutation, (2) there were no mutations outside the v1 loop, and (3) the number of sites mutated once, twice, etc. among these 40 clones approximated a Poisson distribution. When libraries met these criteria, colonies were scraped from all transformation plates and plasmids were directly purified, without further growth to avoid amplification bias, using NucleoBond Xtra midiprep kits (Takara Bio).

### Cell lines

HEK-293T/17 (CVCL_1926) and CRFK (CVCL_2426) cells were grown on tissue-culture treated plates in high-glucose and L-glutamine containing DMEM (Thermo Fisher, Waltham, MA, USA) supplemented with 1x penicillin/streptomycin (Thermo Fisher) and 10% fetal bovine serum (Thermo Fisher). Cell lines were purchased from ATCC (Manassas, VA, USA) and confirmed to be mycoplasma free by MycoProbe kit (R&D Systems, Minneapolis, MN, USA). Cells were grown at 37 °C, 5% CO_2_ in humidified incubators and passaged by digestion with 0.05% trypsin-EDTA (Thermo Fisher). Cell counting was performed using a TC20 automated cell counter (BioRad, Hercules, CA, USA).

### Virus production, titering, and transduction

HEK-293T/17 were seeded at 5 x 10^5^ cells/well in 6-well plates the day prior to transfection. Transfections were performed with Trans-IT 293T transfection reagent (Thermo Fisher) according to manufacturer’s instructions, using 3 μL reagent per μg DNA. All transfected DNA was purified using PureYield mini or NuceloBond midi kits to minimize LPS contamination and quantified by NanoDrop (Thermo Fisher) A260. To produce HIV-1, each well was transfected with 1 μg of p8.9NdSB, 667 ng of pHIV-ZsGreen, and 333 ng of pMD2.G. N-MLV transfections contained 1 μg of pQCXIP-eGFP, 667 ng of pCIG3N, and 333 ng of pMD2.G. HIV-2, SIVcpz, and SIVmac transfections contained 1 μg of pALPS-eGFP, 333 ng of pMD2.G, and 667 ng of either p8.9NdSB HIV-2 CA, pHIV-Gb2, or pHIV-MAC, respectively. TRIM5α-transducing virus was produced using 1 μg of the appropriate TRIM5α construct, 600 ng of JK3, 300 ng of L-VSV-G, and 100 ng of CMV-Tat. After 24 hr, media was replaced with 1 mL of fresh media. Virus was harvested at 48 hr post-transfection. To harvest, media was pelleted at 500 x g, and supernatant was removed, aliquoted, and snap frozen in liquid nitrogen. To increase titers for HIV-2 and N-MLV, and in some cases HIV-1, virus was concentrated prior to freezing. To concentrate, virus was pelleted through a 20% sucrose cushion at 23,000 rpm (~70,000 x g) for 1 hr at 4 °C. Pellets were air dried for 5 min, and then resuspended in fresh media for 24 hr with periodic gentle vortexing.

All viruses were titered under conditions most closely mimicking their large-scale use. CRFK cells were seeded at 1 x 10^5^ cells/mL the day prior to transduction. Freshly thawed viruses were serially diluted and replaced cellular media at ½ x volume. No transducing reagent was used for GFP-marked retroviral VLPs; TRIM5α-transducing VLPs were supplemented with 10 μg/mL polybrene. Plates were centrifuged at 1100 x g for 30 min and then incubated at 37 °C. The following day, virus was removed and cells were fed fresh media, which contained 6 μg/mL puromycin for TRIM5α-transducing VLPs only. For GFP-marked retroviral VLPs, transduction efficiency was monitored by flow cytometry 72 hr after transduction. For TRIM5α- transducing VLPs, cell survival was monitored daily by estimating cell confluence, until untransduced cells were completely dead (no surface-adhered cells). Media was replaced with fresh puromycin-containing media every 2-3 days, and cells were passaged into larger well format as needed. Multiplicity of infection (MOI) for serial dilutions was estimated by Poisson distribution; for example, ~63% of cells are expected to be transduced at least once and thus survive selection with an MOI of 1.

To stably transduce TRIM5α, we chose an MOI of ~0.33 (25-30% survival during titering) to minimize multiple transductions per cell (< 5% probability). CRFK cells were seeded in 6-well plates at 2 x 10^5^ cells/well the day prior to transduction; for deep mutational scanning libraries, sufficient cells were seeded to generate at least 500x independent transductions for each nucleotide variant (32 codons x 13 sites x 500 ÷ 25% survival = 8.3 x 10^5^ cells). Cells were transduced at the appropriate MOI with 10 μg/mL polybrene and spinoculation (1100 x g, 30 min), then underwent 6 μg/mL puromycin selection starting at 24 hr post-transduction and continuing until untransduced controls were completely dead (usually ~7 days). Upon completion of selection, surviving cells were pooled and maintained in 2 μg/mL puromycin. Passages always maintained at least 5 x 10^5^ cells (1000x library coverage) to avoid bottlenecking library diversity.

### Deep mutational scan, sequencing, and enrichment calculation

CRFK cells expressing a TRIM5α deep mutational scanning library were seeded in 12-well plates at 1 x 10^5^ cells/well the day prior to viral infection. Sufficient wells were seeded for at least 1000x library coverage among target cells to be sorted 4 days later (assuming at least 2 doublings in that time, with sorting frequency typically ~5% of cells as estimated beforehand by viral titering against DMS library-expressing cells). Thus, each biological replicate began with at least 2.4 x 10^6^ cells seeded from the same CRFK library.

Libraries were infected with HIV-1-GFP or N-MLV-GFP the following day. For loss-of-restriction experiments (Figures 4-6), we chose viral doses that were restricted by WT TRIM5α to < 1%, as determined during preliminary titering experiments. For gain-of-HIV-1-restriction by human TRIM5α, we chose a viral dose in which WT TRIM5α was infected to ~98%, in order to minimize uninfected GFP-negative cells. Infection efficiency was monitored by parallel infection of controls (empty vector, WT TRIM5α, uninfected negative control). Cells were infected by spinoculation (1100 x g, 30 min) and media was replaced 24 hr post-infection. Infected cells were incubated an additional 48 hr to increase GFP expression levels. Cells were harvested by trypsinization, pelleted, and vigorously resuspended as well as filtered (0.7 μm) to minimize aggregation. Cells were FACS sorted, with stringent gating on size, single cells, and presence or absence of GFP (for loss- or gain-of-restriction, respectively). At least 4 x 10^5^ cells (1000x library coverage) were sorted for each biological replicate. For gain-of-HIV-1-restriction by human TRIM5α, sorted GFP-negative cells were pelleted and re-seeded at 1 x 10^5^ cells/well for a second round of infection, at the same dose, the following day, in order to enrich true restrictors and deplete cells uninfected by chance. Infection, harvest, and FACS sorting were all performed identically, except that apparent HIV-1 restriction by pooled variants was improved in the second round of enrichment (~50% GFP-negative compared to ~10% in the first round of infection). Sorted cells were pelleted, resuspended in PBS, and genomic DNA was harvested using Blood & Cell Culture DNA Mini kits (Qiagen, Hilden, Germany). Input samples were harvested from infected but unsorted cells for each replicate.

Illumina libraries were constructed from genomic DNA by 2-step PCR amplification using Q5 polymerase. The first PCR amplified the v1 loop of TRIM5α and added adapters; the second set of PCR primers annealed to these adapters and added a unique 8 bp i7 Nextera barcode as well as P5 and P7 adapters for flow cell binding (see Table 1). Genomic DNA from each sample (2 input replicates, 2 sorted replicates for each experiment) was amplified in 3 separate PCR tubes, with 500 ng of genomic DNA per tube, to offset random PCR jackpotting. This sampled a total of 1.5 μg of DNA, which represents ~500x library coverage, assuming 6.6 pg gDNA/cell and a single TRIM5α integration/cell. After 15 cycles of amplification, samples were digested for 15 min at 37 °C with 5 μL of ExoI (NEB) to remove first round primers. PCR products were then pooled from triplicate tubes, purified by QIAquick PCR purification kit (Qiagen), and the entire elution was divided between 3 separate PCR tubes for 18 cycles of second round amplification. Barcoded PCR products (234 bp) were pooled from triplicate tubes and purified by double-sided size selection using Ampure beads (Beckman Coulter, Pasadena, CA, USA). In brief, large DNA was removed by incubation with 0.8x bead volume and magnetization; PCR products were bound from the supernatant with 1.5x bead volume, washed with 80% ethanol, and eluted in water. PCR product purity was confirmed by gel electrophoresis. Samples were then pooled at equimolar ratios and Illumina sequenced (MiSeq-v2) with single-end reads. One read was generated using the i7 index primer for the 8 bp barcode, and a second read used a custom sequencing primer, which annealed immediately adjacent to the v1 loop (33 bp read for human, 39 bp for rhesus TRIM5α). PhiX was included at 15% in sequencing runs to increase per-bp-diversity, since the majority (10/11 for human or 12/13 for rhesus) of reads should not randomize any given codon.

Reads counts for each unique nucleotide sequence from all 4 samples in an experiment were compiled into a single tsv file. Sequences that differed from WT by more than 1 codon, or sequences in which codons did not end in C or G, were filtered from the dataset; these largely had only a few reads per sample and represented sequencing errors. Reads counts were normalized to total counts per million (cpm) within each barcoded sample. Sequences with low read counts (< 50 cpm) were excluded as they were found to introduce noise (poor correlation between replicates and across codons). Enrichment was calculated as the ratio of sorted to input cpm. The average and standard deviation of enrichment was calculated across both replicates of all synonymous codons to determine statistics at the amino acid level, except for WT variants, where we show each synonymous variant separately (averaged across replicates) to better visualize WT variance. Amino acid enrichment values were plotted in waterfall plots (descending order of enrichment), scatter plots (comparing replicates), and double-gradient heat maps (comparing amino acid variants at each position, with baseline value [white] set to the average for WT enrichment) using GraphPad Prism. R scripts for data analysis, including all filtering, normalization, and calculations, as well as raw sequence reads have been uploaded to Github: https://github.com/jtenthor/T5DMS_data_analysis.

### Calculation of fold virus inhibition

Viral inhibition by TRIM5α constructs was always compared to CRFK cells transduced with empty vector. CRFK lines were seeded in 96-well plates at 1 x 10^4^ cells/well the day prior to transduction. Media was removed and replaced with serial 3-fold dilutions of the appropriate GFP-marked retrovirus. Serial dilutions were started at titers that yielded ~95% infection in untransduced CRFK. Plates were centrifuged at 1100 x g for 30 min and then incubated at 37 °C. The following day, virus was removed and cells were fed fresh media. Cells were harvested by trypsinization 72 hr after transduction and analyzed by flow cytometry for GFP fluorescence. Cells were gated on size (FSC vs. SSC), single cells (FSC height vs. area), and GFP+ as compared to negative control (FITC vs. PE empty channel).

Fold inhibition was calculated by comparing ID_10_, the amount of virus required to infect 10% of cells, between TRIM5α and empty vector. Infection (% GFP-positive) was plotted against viral dose, both on logarithmic scale, as in Figure 2E. Infection points < 0.5% or > 50% GFP-positive were excluded due to increased noise or curve saturation, respectively, yielding a simple linear relationship. A linear regression (against log-transformed data) was then used to calculate the viral dose corresponding to 10% infection (back-calculated to linear scale), and the dose for TRIM5α was divided by that for empty vector. This method was used to calculate fold inhibition for all viruses except human TRIM5α against N-MLV, as we could not consistently achieve infection greater than 1% for WT human TRIM5α; we therefore report raw infection data. All fold inhibition was calculated from at least 3 independent experiments, which were performed either in biological singlicate or duplicate.

### Immunoblot

CRFK cells stably expressing TRIM5α-HA variants were harvested by trypsinization, washed in PBS, and counted; 10^6^ cells were lysed for 15 min on ice in 100 μL pre-chilled lysis buffer (50 mM Tris, pH 8, 150 mM NaCl, 1% Triton-X100, 1x cOmplete EDTA-free protease inhibitor cocktail [Roche, Basel, Switzerland]). Lysates were pelleted at 20,000 x g for 15 min at 4 °C. Supernatants were quantified by Bradford protein assay (BioRad) and normalized to load equal protein across all samples (usually 10-25 μg per lane). Samples were boiled for 5 min in Laemmli Sample Buffer (BioRad) supplemented with 5% β-mercaptoethanol and loaded onto Mini-PROTEAN TGX stain-free gels (BioRad). Gels were run in Tris/Glycine/SDS buffer (BioRad) for 50 min at 150 V, then transferred semi-dry for 7 min at 1.3 mV using Trans-Blot Turbo 0.2 μm nitrocellulose transfer packs and the Trans-Blot Turbo transfer system (BioRad). Blots were blocked with Odyssey blocking buffer (LI-COR, Lincoln, NE, USA), then probed with mouse anti-HA at 1:1000 (AB_2565335, Biolegend, San Diego, CA, USA) and rabbit anti-β-actin at 1:5000 (AB_2305186, Abcam, Cambridge, UK). All antibodies were diluted in TBST with 5% bovine serum albumin (Sigma Aldrich, St. Louis, MO, USA). Blots were washed in TBST and probed with IRDye 680RD donkey anti-mouse (AB_10953628, LI-COR) and IRDye 800CW donkey anti-rabbit (AB_621848, LI-COR), both diluted 1:10000. Blots were washed and scanned at 680 and 800 nm. HA intensities were quantified using ImageJ and normalized to actin, then compared to WT TRIM5α to determine relative expression levels.

### TRIM5α phylogeny, rapid evolution analysis, and evolutionary accessibility

A tBLASTn search of human TRIM5α (NP_149023.2) against primate genomes returned 29 unique simian primate orthologs of TRIM5α. We excluded New World monkey sequences as they share a 9-amino acid deletion in the v1 loop. Open reading frames of the following sequences were translation aligned using MUSCLE: human (*Homo sapiens*, NM_033034.2), chimpanzee (*Pan troglodytes*, NM_001012650.1), bonobo (*Pan paniscus*, XM_003819046.3), gorilla (*Gorilla gorilla*, NM_001279549.1), Sumatran orangutan (*Pongo abelii*, NM_001131070.1), Bornean orangutan (*Pongo pygmaeus*, AY923179.2), white-handed gibbon (*Hylobates lar*, AY923180.1), white-cheeked gibbon (*Nomascus leucogenys*, NM_001280113.1), crab-eating macaque (*Macaca fascicularis*, NM_001283295.1), rhesus macaque (*Macaca mulatta*, NM_001032910.1), olive baboon (*Papio anubis*, NM_001112632.1), collared mangabey (*Cercocebus torquatus*, KP743974.1), sooty mangabey (*Cercocebus atys*, NM_001305964.1), drill (*Mandrillus leucophaeus*, XM_011971974.1), Wolf’s guenon (*Cercopithecus wolfi*, KP743973.1), red guenon (*Erythrocebus patas*, AY740619.1), grivet (*Cercopithecus aethiops*, AY669399.1), tantalus monkey (*Chlorocebus tantalus*, AY740613.1), African green monkey (*Chlorocebus sabaeus*, XM_008019877.1), vervet monkey (*Chlorocebus pygerythrus*, AY740612.1), golden snub-nosed monkey (*Rhinopithecus roxellana*, XM_010364548.1), and Angola colobus (*Colobus angolensis*, XM_011963593.1).

A PHYML tree was built using the HKY85 substitution model with 100 bootstraps and rooted on human TRIM6 (NM_001003818.3). The unrooted tree was used for site-specific PAML analysis (Z. Yang, 1997) using both F3×4 and F61 codon models to ensure robust results. We performed maximum likelihood (ML) tests comparing model 7 (neutral selection beta distribution) to model 8 (beta distribution with positive selection allowed). In each case, the model allowing positive selection gave the best fit to the data (p < 0.0001, chi-squared test on 2x ΔML with 2 df). Model 8 also identified rapidly evolving sites with a Bayes Empirical Bayes posterior probability > 0.95. We report residues that meet this threshold for rapid evolution under both the F3×4 and F61 codon models.

Evolutionarily accessible amino acids were defined as 1 nucleotide substitution away from the wild-type sequence. For Figure 1B, we determined all amino acids that were accessible from any of the 22 aligned sequences, then determined the fraction of these amino acids that were represented in our alignment.

## ACKNOWLEDGEMENTS

We thank all members of the Malik and Emerman labs, especially Shirleen Soh and Molly OhAinle, for feedback and advice. We thank Tera Levin, Kevin Forsberg, Molly Ohainle, Tyler Starr, Russell Vance, Patrick Mitchell, Janet Young, Nicholas Chesarino, Pravrutha Raman, Phoebe Hsieh, Shirleen Soh, and Rick McLaughlin for comments on the manuscript. The Fred Hutchinson Genomics Core performed Illumina sequencing. Chimeric HIV-1 gag/pol vectors with SIV CA were a gift from Theodora Hatziioannou. HIV-1 and HIV-2 gag/pol vectors were a gift from Jeremy Luban. Work was supported by the Hanna H. Gray fellowship (J.L.T.), an EXROP award (A.S.), and an Investigator award (H.S.M.) from HHMI, in addition to grants from the Mathers Foundation and NIAID (HARC [HIV Accessory and Regulatory Complexes] center, P50 AI082250, PI Nevan Krogan, subaward to M.E., H.S.M.). The authors declare no competing interests.

## AUTHOR CONTRIBUTIONS

J.L.T., M.E., and H.S.M. conceived the study, designed experiments, and wrote the manuscript. J.L.T., C.Y., and A.S. performed and analyzed experiments. C.Y. edited the manuscript.

## Notes

https://github.com/jtenthor/T5DMS_data_analysis

### Competing Interest Statement

The authors have declared no competing interest.

## REFERENCES

Berthoux, L., Sebastian, S., Sokolskaja, E., & Luban, J. (2004). Lv1 Inhibition of Human Immunodeficiency Virus Type 1 Is Counteracted by Factors That Stimulate Synthesis or Nuclear Translocation of Viral cDNA. Journal of Virology, 78(21), 11739–11750. http://doi.org/10.1128/JVI.78.21.11739-11750.2004

Biris, N., Yang, Y., Taylor, A. B., Tomashevski, A., Guo, M., Hart, P. J., et al. (2012). Structure of the rhesus monkey TRIM5α PRYSPRY domain, the HIV capsid recognition module. Proceedings of the National Academy of Sciences of the United States of America, 109(33), 13278–13283. http://doi.org/10.1073/pnas.1203536109

Bock, M., Bishop, K. N., Towers, G., & Stoye, J. P. (2000). Use of a transient assay for studying the genetic determinants of Fv1 restriction. Journal of Virology, 74(16), 7422–7430. http://doi.org/10.1128/jvi.74.16.7422-7430.2000

Clarke, L., Fairley, S., Zheng-Bradley, X., Streeter, I., Perry, E., Lowy, E., et al. (2017). The international Genome sample resource (IGSR): A worldwide collection of genome variation incorporating the 1000 Genomes Project data. Nucleic Acids Research, 45(D1), D854–D859. http://doi.org/10.1093/nar/gkw829

Colón-Thillet, R., Hsieh, E., Graf, L., McLaughlin, R. N., Young, J. M., Kochs, G., et al. (2019). Combinatorial mutagenesis of rapidly evolving residues yields super-restrictor antiviral proteins. PLoS Biology, 17(10), e3000181–18. http://doi.org/10.1371/journal.pbio.3000181

Compton, A. A., & Emerman, M. (2013). Convergence and divergence in the evolution of the APOBEC3G-Vif interaction reveal ancient origins of simian immunodeficiency viruses. PLoS Pathogens, 9(1), e1003135. http://doi.org/10.1371/journal.ppat.1003135

Daugherty, M. D., & Malik, H. S. (2012). Rules of Engagement: Molecular Insights from Host-Virus Arms Races. Annual Review of Genetics, 46(1), 677–700. http://doi.org/10.1146/annurev-genet-110711-155522

Daugherty, P. S., Chen, G., Iverson, B. L., & Georgiou, G. (2000). Quantitative analysis of the effect of the mutation frequency on the affinity maturation of single chain Fv antibodies. Proceedings of the National Academy of Sciences of the United States of America, 97(5), 2029–2034. http://doi.org/10.1073/pnas.030527597

Draghi, J. A., Parsons, T. L., Wagner, G. P., & Plotkin, J. B. (2010). Mutational robustness can facilitate adaptation. Nature, 1–3. http://doi.org/10.1038/nature08694

Duggal, N. K., & Emerman, M. (2012). Evolutionary conflicts between viruses and restriction factors shape immunity. Nature Reviews Immunology, 1–9. http://doi.org/10.1038/nri3295

Fowler, D. M., Araya, C. L., Fleishman, S. J., Kellogg, E. H., Stephany, J. J., Baker, D., & Fields, S. (2010). High-resolution mapping of protein sequence-function relationships. Nature Methods, 7(9), 741–746. http://doi.org/10.1038/nmeth.1492

Guo, H. H., Choe, J., & Loeb, L. A. (2004). Protein tolerance to random amino acid change. Proceedings of the National Academy of Sciences of the United States of America, 101(25), 9205–9210. http://doi.org/10.1073/pnas.0403255101

Hayden, E. J., Ferrada, E., & Wagner, A. (2011). Cryptic genetic variation promotes rapid evolutionary adaptation in an RNA enzyme. Nature, 474(7349), 92–95. http://doi.org/10.1038/nature10083

Jimenez-Guardeño, J. M., Apolonia, L., Betancor, G., & Malim, M. H. (2019). Immunoproteasome activation enables human TRIM5α restriction of HIV-1. Nature Microbiology, 1–11. http://doi.org/10.1038/s41564-019-0402-0

Kim, K., Dauphin, A., Komurlu, S., McCauley, S. M., Yurkovetskiy, L., Carbone, C., et al. (2019). Cyclophilin A protects HIV-1 from restriction by human TRIM5α. Nature Microbiology, 1–17. http://doi.org/10.1038/s41564-019-0592-5

Kirmaier, A., Wu, F., Newman, R. M., Hall, L. R., Morgan, J. S., O’Connor, S., et al. (2010). TRIM5 Suppresses Cross-Species Transmission of a Primate Immunodeficiency Virus and Selects for Emergence of Resistant Variants in the New Species. PLoS Biology, 8(8), e1000462–12. http://doi.org/10.1371/journal.pbio.1000462

Kratovac, Z., Virgen, C. A., Bibollet-Ruche, F., Hahn, B. H., Bieniasz, P. D., & Hatziioannou, T. (2008). Primate lentivirus capsid sensitivity to TRIM5 proteins. Journal of Virology, 82(13), 6772–6777. http://doi.org/10.1128/JVI.00410-08

Lascano, J., Uchil, P. D., Mothes, W., & Luban, J. (2015). TRIM5 Retroviral Restriction Activity Correlates with the Ability To Induce Innate Immune Signaling. Journal of Virology, 90(1), 308–316. http://doi.org/10.1128/JVI.02496-15

Li, Y., Li, X., Stremlau, M., Lee, M., & Sodroski, J. (2006). Removal of arginine 332 allows human TRIM5αlpha to bind human immunodeficiency virus capsids and to restrict infection. Journal of Virology, 80(14), 6738–6744. http://doi.org/10.1128/JVI.00270-06

Li, Y.-L., Chandrasekaran, V., Carter, S. D., Woodward, C. L., Christensen, D. E., Dryden, K. A., et al. (2016). Primate TRIM5 proteins form hexagonal nets on HIV-1 capsids. eLife, 5. http://doi.org/10.7554/eLife.16269

Maillard, P. V., Reynard, S., Serhan, F., Turelli, P., & Trono, D. (2007). Interfering residues narrow the spectrum of MLV restriction by human TRIM5αlpha. PLoS Pathogens, 3(12), e200. http://doi.org/10.1371/journal.ppat.0030200

McCarthy, K. R., Kirmaier, A., Autissier, P., & Johnson, W. E. (2015). Evolutionary and Functional Analysis of Old World Primate TRIM5 Reveals the Ancient Emergence of Primate Lentiviruses and Convergent Evolution Targeting a Conserved Capsid Interface. PLoS Pathogens, 11(8), e1005085. http://doi.org/10.1371/journal.ppat.1005085

McCauley, S. M., Kim, K., Nowosielska, A., Dauphin, A., Yurkovetskiy, L., Diehl, W. E., & Luban, J. (2018). Intron-containing RNA from the HIV-1 provirus activates type I interferon and inflammatory cytokines. Nature Communications, 1–10. http://doi.org/10.1038/s41467-018-07753-2

McEwan, W. A., Schaller, T., Ylinen, L. M., Hosie, M. J., Towers, G. J., & Willett, B. J. (2009). Truncation of TRIM5 in the Feliformia Explains the Absence of Retroviral Restriction in Cells of the Domestic Cat. Journal of Virology, 83(16), 8270–8275. http://doi.org/10.1128/JVI.00670-09

McLaughlin, R. N., Poelwijk, F. J., Raman, A., Gosal, W. S., & Ranganathan, R. (2012). The spatial architecture of protein function and adaptation. Nature, 491(7422), 138–142. http://doi.org/10.1038/nature11500

Mitchell, P. S., Patzina, C., Emerman, M., Haller, O., Malik, H. S., & Kochs, G. (2012). Evolution-Guided Identification of Antiviral Specificity Determinants in the Broadly Acting Interferon-Induced Innate Immunity Factor MxA. Cell Host and Microbe, 12(4), 598–604. http://doi.org/10.1016/j.chom.2012.09.005

Newman, R. M., Hall, L., Connole, M., Chen, G.-L., Sato, S., Yuste, E., et al. (2006). Balancing selection and the evolution of functional polymorphism in Old World monkey TRIM5αlpha. Proceedings of the National Academy of Sciences of the United States of America, 103(50), 19134–19139. http://doi.org/10.1073/pnas.0605838103

OhAinle, M., Helms, L., Vermeire, J., Roesch, F., Humes, D., Basom, R., et al. (2018). A virus-packageable CRISPR screen identifies host factors mediating interferon inhibition of HIV. eLife, 7, 783. http://doi.org/10.7554/eLife.39823

Ohkura, S., Yap, M. W., Sheldon, T., & Stoye, J. P. (2006). All three variable regions of the TRIM5αlpha B30.2 domain can contribute to the specificity of retrovirus restriction. Journal of Virology, 80(17), 8554–8565. http://doi.org/10.1128/JVI.00688-06

Owens, C. M., Yang, P. C., Gottlinger, H., & Sodroski, J. (2003). Human and Simian Immunodeficiency Virus Capsid Proteins Are Major Viral Determinants of Early, Postentry Replication Blocks in Simian Cells. Journal of Virology, 77(1), 726–731. http://doi.org/10.1128/JVI.77.1.726-731.2003

Perron, M. J., Stremlau, M., & Sodroski, J. (2006). Two surface-exposed elements of the B30.2/SPRY domain as potency determinants of N-tropic murine leukemia virus restriction by human TRIM5αlpha. Journal of Virology, 80(11), 5631–5636. http://doi.org/10.1128/JVI.00219-06

Pertel, T., Hausmann, S., Morger, D., Züger, S., Guerra, J., Lascano, J., et al. (2011). TRIM5 is an innate immune sensor for the retrovirus capsid lattice. Nature, 472(7343), 361–365. http://doi.org/10.1038/nature09976

Pham, Q. T., Bouchard, A., Grütter, M. G., & Berthoux, L. (2010). Generation of human TRIM5αlpha mutants with high HIV-1 restriction activity. Gene Therapy, 17(7), 859–871. http://doi.org/10.1038/gt.2010.40

Pham, Q. T., Veillette, M., Brandariz-Nuñez, A., Pawlica, P., Thibert-Lefebvre, C., Chandonnet, N., et al. (2013). A novel aminoacid determinant of HIV-1 restriction in the TRIM5α variable 1 region isolated in a random mutagenic screen. Virus Research, 173(2), 306–314. http://doi.org/10.1016/j.virusres.2013.01.013

Pizzato, M., McCauley, S. M., Neagu, M. R., Pertel, T., Firrito, C., Ziglio, S., et al. (2015). Lv4 Is a Capsid-Specific Antiviral Activity in Human Blood Cells That Restricts Viruses of the SIVMAC/SIVSM/HIV-2 Lineage Prior to Integration. PLoS Pathogens, 11(7), e1005050–29. http://doi.org/10.1371/journal.ppat.1005050

Sawyer, S. L., Emerman, M., & Malik, H. S. (2004). Ancient Adaptive Evolution of the Primate Antiviral DNA-Editing Enzyme APOBEC3G. PLoS Biology, 2(9), e275–8. http://doi.org/10.1371/journal.pbio.0020275

Sawyer, S. L., Wu, L. I., Emerman, M., & Malik, H. S. (2005). Positive selection of primate TRIM5αlpha identifies a critical species-specific retroviral restriction domain. Proceedings of the National Academy of Sciences of the United States of America, 102(8), 2832–2837. http://doi.org/10.1073/pnas.0409853102

Sebastian, S., & Luban, J. (2005). TRIM5αlpha selectively binds a restriction-sensitive retroviral capsid. Retrovirology, 2, 40. http://doi.org/10.1186/1742-4690-2-40

Selyutina, A., Persaud, M., Simons, L. M., Bulnes-Ramos, A., Buffone, C., Martinez-Lopez, A., et al. (2020). Cyclophilin A Prevents HIV-1 Restriction in Lymphocytes by Blocking Human TRIM5&#x03B1; Binding to the Viral Core. Cell Reports, 30(11), 3766–3777.e6. http://doi.org/10.1016/j.celrep.2020.02.100

Sheng, Z., Schramm, C. A., Kong, R., NISC Comparative Sequencing Program, Mullikin, J. C., Mascola, J. R., et al. (2017). Gene-Specific Substitution Profiles Describe the Types and Frequencies of Amino Acid Changes during Antibody Somatic Hypermutation. Frontiers in Immunology, 8, 537. http://doi.org/10.3389/fimmu.2017.00537

Smith, J. M. (1970). Natural selection and the concept of a protein space. Nature, 225(5232), 563–564. http://doi.org/10.1038/225563a0

Starr, T. N., Picton, L. K., & Thornton, J. W. (2017). Alternative evolutionary histories in the sequence space of an ancient protein. Nature, 549(7672), 409–413. http://doi.org/10.1038/nature23902

Stiffler, M. A., Hekstra, D. R., & Ranganathan, R. (2015). Evolvability as a Function of Purifying Selection in TEM-1 β-Lactamase. Cell, 160(5), 882–892. http://doi.org/10.1016/j.cell.2015.01.035

Stremlau, M., Owens, C. M., Perron, M. J., Kiessling, M., Autissier, P., & Sodroski, J. (2004). The cytoplasmic body component TRIM5αlpha restricts HIV-1 infection in Old World monkeys. Nature, 427(6977), 848–853. http://doi.org/10.1038/nature02343

Stremlau, M., Perron, M., Lee, M., Li, Y., Song, B., Javanbakht, H., et al. (2006). Specific recognition and accelerated uncoating of retroviral capsids by the TRIM5αlpha restriction factor. Proceedings of the National Academy of Sciences of the United States of America, 103(14), 5514–5519. http://doi.org/10.1073/pnas.0509996103

Stremlau, M., Perron, M., Welikala, S., & Sodroski, J. (2005). Species-specific variation in the B30.2(SPRY) domain of TRIM5αlpha determines the potency of human immunodeficiency virus restriction. Journal of Virology, 79(5), 3139–3145. http://doi.org/10.1128/JVI.79.5.3139-3145.2005

Suckow, J., Markiewicz, P., Kleina, L. G., Miller, J., Kisters-Woike, B., & Müller-Hill, B. (1996). Genetic studies of the Lac repressor. XV: 4000 single amino acid substitutions and analysis of the resulting phenotypes on the basis of the protein structure. Journal of Molecular Biology, 261(4), 509–523. http://doi.org/10.1006/jmbi.1996.0479

Tareen, S. U., & Emerman, M. (2011). Human Trim5α has additional activities that are uncoupled from retroviral capsid recognition. Virology, 409(1), 113–120. http://doi.org/10.1016/j.virol.2010.09.018

Van Valen, L. (1973). A new evolutionary law. Evol. Theory, (1), 1–30.

Veillette, M., Bichel, K., Pawlica, P., Freund, S. M. V., Plourde, M. B., Pham, Q. T., et al. (2013). The V86M mutation in HIV-1 capsid confers resistance to TRIM5α by abrogation of cyclophilin A-dependent restriction and enhancement of viral nuclear import. Retrovirology, 10(1), 1–14. http://doi.org/10.1186/1742-4690-10-25

Welm, B. E., Dijkgraaf, G. J. P., Bledau, A. S., Welm, A. L., & Werb, Z. (2008). Lentiviral Transduction of Mammary Stem Cells for Analysis of Gene Function during Development and Cancer. Cell Stem Cell, 2(1), 90–102. http://doi.org/10.1016/j.stem.2007.10.002

Wu, F., Kirmaier, A., Goeken, R., Ourmanov, I., Hall, L., Morgan, J. S., et al. (2013). TRIM5 alpha drives SIVsmm evolution in rhesus macaques. PLoS Pathogens, 9(8), e1003577. http://doi.org/10.1371/journal.ppat.1003577

Yang, Z. (1997). PAML: a program package for phylogenetic analysis by maximum likelihood. Computer Applications in the Biosciences: CABIOS, 13(5), 555–556. http://doi.org/10.1093/bioinformatics/13.5.555

Yap, M. W., Nisole, S., & Stoye, J. P. (2005). A single amino acid change in the SPRY domain of human Trim5alpha leads to HIV-1 restriction. Current Biology: CB, 15(1), 73–78. http://doi.org/10.1016/j.cub.2004.12.042

